# Acculturative orientations among Hispanic/Latinx caregivers in the ABCD Study: Associations with caregiver and youth mental health and youth brain function

**DOI:** 10.1101/2022.07.24.501248

**Authors:** Alan Meca, Julio A. Peraza, Michael C. Riedel, Willie Hale, Jeremy W. Pettit, Erica D. Musser, Taylor Salo, Jessica S. Flannery, Katherine L. Bottenhorn, Anthony S. Dick, Rosario Pintos Lobo, Laura M. Ucros, Chelsea A. Greaves, Samuel W. Hawes, Mariana Sanchez, Marybel R. Gonzalez, Matthew T. Sutherland, Raul Gonzalez, Angela R. Laird

## Abstract

**Background:** Population-based neuroscience offers opportunities to examine important but understudied sociocultural factors, such as acculturation. Acculturation refers to the extent to which an individual retains their cultural heritage and / or adopts the receiving society’s culture and is particularly salient among Hispanic/Latinx immigrants. Specific acculturative orientations have been linked to vulnerability to substance use, depression, and suicide and are known to influence family dynamics between caregivers and their children.

**Methods:** We investigated first- and second-generation Hispanic/Latinx caregivers in the ABCD Study and examined how caregivers’ acculturative orientation impacts their mental health, as well as the mental health of their children. In addition, we evaluated how caregiver orientation is associated with adolescent socio-affiliative neural function in the ventromedial prefrontal cortex, insula, and temporoparietal junction.

**Results:** We identified two caregiver acculturation profiles: bicultural (retains heritage culture while adopting US culture) and detached (discards heritage culture and rejects US culture). Bicultural caregivers exhibited fewer symptoms of depression, avoidant behaviors, and inattention compared to detached caregivers; further, youth exhibited similar internalizing effects across caregiver profiles. Moreover, youth with bicultural caregivers displayed increased resting-state brain activity in the left insula; however, differences in long-range functional connectivity were not significant.

**Conclusions:** Caregiver acculturation is an important familial-environmental factor in Hispanic/Latinx families linked to significant differences in caregiver and youth mental health and youth insula activity. Future work should examine sociocultural and neurodevelopmental changes across adolescence to assess health outcomes and determine whether localized, corticolimbic brain effects are ultimately translated into long-range connectivity differences.

## INTRODUCTION

Acculturation broadly refers to the extent to which an individual retains their cultural heritage and / or adopts the receiving society’s culture (1), and has been hypothesized to play an important role in accounting for health disparities, particularly among Hispanic/Latinx populations (2). Within the United States (US), Hispanic/Latinx people are among the largest and fastest-growing immigrant groups (3) and acculturative processes are salient among first- and second-generation, and to some extent, third-generation, Hispanic/Latinx immigrants (1). Contemporary views of acculturation draw on Berry’s model (4), which casts acculturation as a bidimensional process consisting of receiving/US culture acquisition and heritage culture retention. This bidimensional conceptualization proposes that individuals can acquire receiving culture without discarding their heritage culture (1). Consistently, research has extensively supported a bidimensional model of acculturation, consisting of distinct heritage and US cultural orientations (1,5–7). Additionally, Berry’s model (8,9) proposes four distinct acculturative orientations: (1) **bicultural** (i.e., acquires the receiving culture and retains the heritage culture); (2) **assimilated** (i.e., acquires the receiving culture and discards the heritage culture); (3) **separated** (i.e., rejects the receiving culture and retains the heritage culture); and (4) **detached** (i.e., rejects the receiving culture and discards the heritage culture)^1^ (**Fig. 1**).

**Figure 1.**
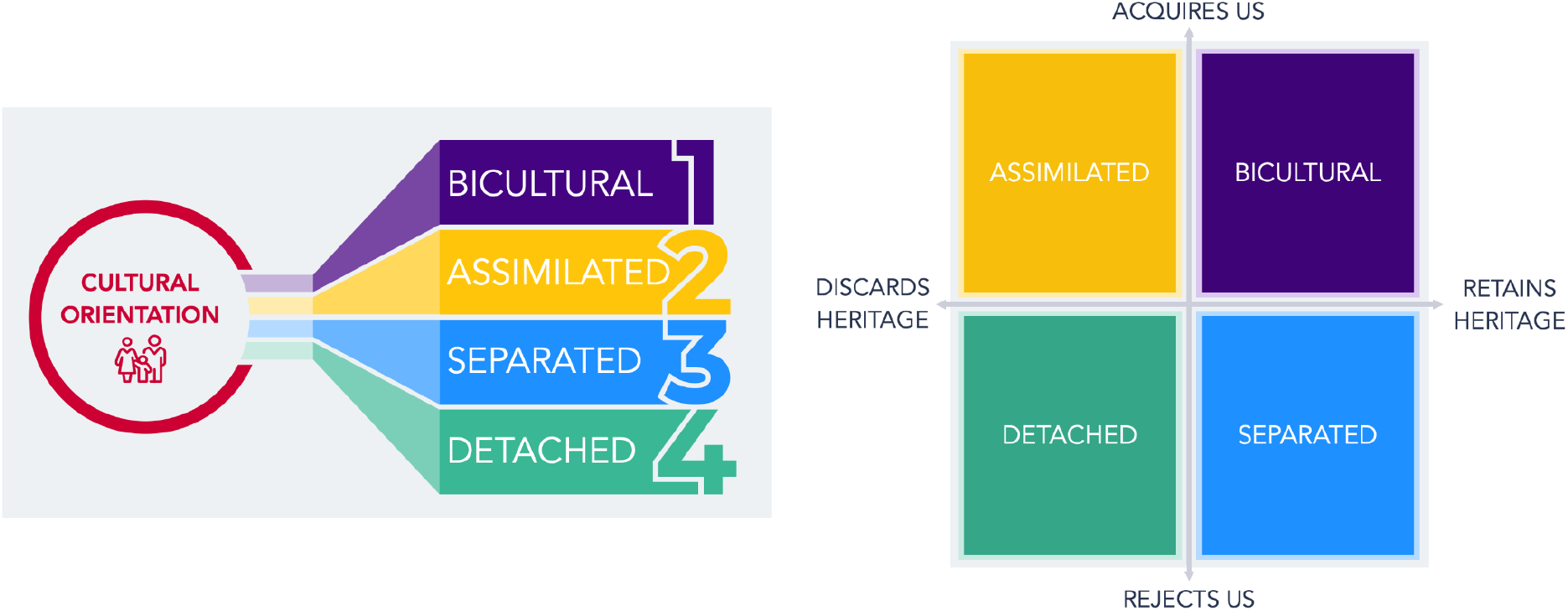
Acculturative Orientation Profiles. Berry’s model proposes four distinct acculturative orientations. Among Hispanic/Latinx immigrants in the US, these profiles include: (1) **bicultural** (i.e., acquires US culture and retains the heritage culture), (2) **assimilated** (i.e., acquires US culture and discards the heritage culture), (3) **separated** (i.e., rejects US culture and retains the heritage culture), and (4) **detached** (i.e., rejects US culture and discards the heritage culture).

A number of studies (6,10–15) have provided mixed support for Berry’s model (4) using person-centered methods, such as latent profile analysis (LPA), that empirically derive acculturation groupings. Among recently immigrated Hispanic/Latinx caregivers, detached and bicultural profiles were identified across heritage and US cultural practices and identity (16). In contrast, three profiles were recently identified among a community sample of undocumented Hispanic/Latinx immigrants: bicultural, detached, and separated (13), while another study identified separated, partially separated, bicultural, and detached profiles (17). Although specific configurations have been inconsistently observed across studies, biculturalism has been consistently identified and characterized as the most adaptive orientation (10,18,19). Indeed, individuals who are bicultural have been found in comparison to other orientations to exhibit fewer symptoms of depression (17,20) and anxiety (20), lower substance use (12), and highest levels of flourishing and life satisfaction (21).

Despite extensive research among Hispanic/Latinx adults, few studies have addressed how acculturation may be related to adolescent (mal)adaptive development and behavior. Adolescence is a critical developmental stage for identity development (22) during which youth establish a self identity, as well as a social identity based on group membership. This significant neurodevelopmental period (23,24) is marked by changes in self-referential cognition (25) that are accompanied by maturational shifts in brain structure and function that occur as a consequence of experiences within one’s social environment (26,27). Such shifts occur in the ventromedial prefrontal cortex (vmPFC), insula, and temporoparietal junction (TPJ) (28), which have been linked to self- and affiliation-based social processes (29). In addition to developing a general sense of self and identity, which is a normative developmental task for all adolescents, Hispanic/Latinx youth are also tasked with establishing an acculturative orientation. Hispanic/Latinx adolescents experience shifts in acculturation, encompassing dynamic changes in values, beliefs, and practices that together reflect their cultural group affiliation. These processes impact one’s sense of self and can be broadly shaped by their caregivers’ own acculturative orientation (30). Indeed, caregivers’ acculturative orientation impacts the degree to which they socialize their children to gravitate towards (or away) from the receiving and heritage cultures (30–32). Thus, it is likely that caregiver acculturation influences not only caregiver mental health, but also has a direct downstream effect on youth mental health. That is, given the strong influence that caregivers have on child and adolescent behavior and psychopathology (33–35), we hypothesize that caregiver biculturalism is associated with fewer mental health problems among their children. Further, drawing on a conceptual framework of adolescent neurobiological susceptibility to social contexts via caregivers and peers (28), we also hypothesize that caregiver acculturation is directly associated with adolescent socio-affiliative neural function.

The overall objective of the current study was to test the hypotheses that caregiver acculturation is associated with youth mental health and brain function. To this end, we first sought to evaluate the acculturative orientations of first- and second-generation Hispanic/Latinx immigrant caregivers in the Adolescent Brain Cognitive Development^SM^ Study (ABCD Study®) (36). We focused on ABCD data acquired at baseline (i.e., 9-10-year-old youth) to establish an understanding of caregiver acculturation among ABCD Hispanic/Latinx families as an important familial influence at the onset of adolescence. To this end, and building on prior work among adult populations (11,13), we aimed to further empirically validate Berry’s model of acculturation by identifying data-driven groupings of ABCD caregivers’ heritage and US cultural orientation at baseline using latent profile analysis (LPA). Consistent with recommended best practices (37), a data-driven approach is critical as it allows researchers to avoid the utilization of arbitrary cut-off thresholds or the inaccurate identification of much-debated acculturation profiles that may not exist within a given sample (38). Once we identified acculturative orientation profiles, we investigated associations with caregiver and youth mental health and youth resting state functional magnetic resonance imaging (rs-fMRI) signatures in self- and affiliation-related circuits. Given the large and demographically diverse ABCD sample (39), and in line with current theory (8–10) and informed by prior results among adult populations (6,10–15,40), we hypothesized that LPA would reveal four acculturation profiles (i.e., bicultural, assimilated, separated, and detached profiles). We further expected that biculturalism would be associated with fewer mental health problems among caregivers and their children. We leveraged prior meta-analytic results to guide analyses of brain regions associated with self- and affiliation-based social processing, including the vmPFC, insula, and TPJ (29). Finally, we discuss additional research needed to more fully understand neurobiological mechanisms linked with acculturative experiences.

## METHODS

### Participants

Participants were selected from the ABCD Study, the largest longitudinal study of brain development and child health in the US (36). Approximately 11,800 youth aged 9.00 to 10.99 years old were enrolled in the ABCD Study across 21 sites in the US (41). Participants within the ABCD Study were recruited through geographically, demographically, and socioeconomically diverse school systems using epidemiologically informed methods to enroll a population-based, demographically diverse sample (39). Data from the ABCD Study are made available by the NIMH Data Archive and the current study utilized data from the ABCD Curated Annual Release 3.0.

### Measures

#### Caregiver-Reported Demographics

Demographic information was provided at baseline by a child’s caregiver, including the child’s age, gender, ethnicity, race, as well as the caregiver’s age, identity, gender, ethnicity, race, education, and combined family income. In addition, caregivers reported the nativity (i.e., country of origin) for the child, parent / guardian, and grandparents (42).

#### Caregiver-Reported Acculturation

Within the robust culture and environment battery at baseline (43), caregivers completed the Vancouver Index of Acculturation (VIA) in English or Spanish. The VIA is a 16-item bidimensional measure with subscales that separately measure heritage and US acculturation (44). Items addressed a range of topics, including traditions, social activities, media, cultural values, and behavioral preferences (**Table S1**). Caregivers were asked to provide their heritage culture as an open-ended item with specific examples provided as prompts (e.g., *“Asian”, “Black/African American”, “Hispanic”, “Native American”, “Jewish”);* those who did not identify a heritage culture were told not to complete the VIA. Initial assessment of VIA data at ABCD baseline indicated high internal consistency across the heritage (α=0.92) and US (α=0.90) subscales, with early data suggesting higher VIA subscale scores for both heritage and US cultures among families at lower risk compared to those at higher risk for adolescent substance use (43).

#### Caregiver and Youth Measures of Mental Health

In addition to measures of demographics and acculturation, the ABCD baseline mental health battery included measures of caregiver and youth mental health (42). Caregivers completed the Adult Self Report (ASR), which assesses behavioral dimensions relevant to adult psychopathology (45) and the Child Behavior Checklist (CBCL) (45), which assesses behavioral dimensions relevant to child psychopathology. NDA provides the age- and gender-normed syndrome and DSM-oriented scale scores of the ASR and CBCL; the DSM-oriented scoring was used in the present study.

### Hispanic/Latinx Sample Selection

A total of 11,878 ABCD participants were recruited at baseline. Data for the present analyses were downloaded from NDA for 2,411 participants who completed their baseline assessment for the ABCD Study and responded *“Yes”* to *“Do you consider the child Hispanic/Latino/Latina?”.* We filtered participant datasets to include only caregivers who: (1) completed the VIA, (2) referenced Hispanic/Latinx culture when completing the VIA, (3) and were either first- or second-generation immigrants from Latin America. The subsequent sample thus consisted of 1,057 caregivers (Mean_Age_=38.31 years, *SD*=6.64 years; 90.4% mothers and 9.6% fathers, 70.5% foreign born) and 1,158 children (52.7% male, 91.7% US born) (**Table 1**). Additional details about the sample are provided in the Supplemental Information (SI).

**Table 1.** Demographic Characteristics.

| **Caregiver (N = 1,057)** |  |  |  |  |  |
| --- | --- | --- | --- | --- | --- |
| **Caregiver Identity** | ***n* (%) or *M* (*SD*)** | **Caregiver Country of Origin** | ***n* (%)** | **ABCD Site** | ***n* (%)** |
| Biological mother | 955 (90.4%) | Argentina | 9 (0.9%) | CHLA | 162 (15.3%) |
| Biological father | 102 (9.6%) | Bolivia | 2 (0.2%) | CUB | 16 (1.5%) |
| **Caregiver Age** | 38.31 yrs (6.64 yrs) | Brazil | 17 (1.6%) | FIU | 293 (27.7%) |
| **Caregiver Nativity** |  | Chile | 5 (0.5%) | LIBR | 44 (4.2%) |
| US born | 312 (29.5%) | Colombia | 39 (3.7%) | MUSC | 2 (0.2%) |
| Foreign born | 745 (70.5%) | Costa Rica | 8 (0.8%) | OHSU | 29 (2.7%) |
| **Caregiver Education** |  | Cuba | 59 (5.6%) | ROC | 3 (0.3%) |
| < High school diploma | 239 (22.6%) | Dominican Republic | 12 (1.1%) | SRI | 43 (4.1%) |
| High school diploma/GED | 167 (15.8%) | Ecuador | 13 (1.2%) | UCLA | 87 (8.2%) |
| Some college | 352 (33.3%) | El Salvador | 27 (2.6%) | UCSD | 249 (23.6%) |
| Bachelor's degree | 172 (16.3%) | Guatemala | 27 (2.6%) | UFL | 13 (1.2%) |
| Postgraduate degree | 123 (11.6%) | Honduras | 26 (2.5%) | UMB | 12 (1.1%) |
| No answer | 4 (0.4%) | Mexico | 406 (38.4%) | UMICH | 17 (1.6%) |
| **Household Income per Year (\$US)** | | Nicaragua | 29 (2.7%) | UMN | 6 (0.6%) |
| <\$5,000 | 49 (4.6%) | Panama | 2 (0.2%) | UPMC | 1 (0.1%) |
| \$5,000-\$11,999 | 65 (6.1%) | Paraguay | 1 (0.1%) | UTAH | 35 (3.3%) |
| \$12,000-\$15,999 | 53 (5.0%) | Peru | 27 (2.6%) | UVM | 4 (0.4%) |
| \$16,000-\$24,999 | 103 (9.7%) | Uruguay | 3 (0.3%) | UWM | 2 (0.2%) |
| \$25,000-\$34,999 | 126 (11.9%) | United States | 312 (29.5%) | VCU | 11 (1.0%) |
| \$35,000-\$49,999 | 137 (13.0%) | Venezuela | 33 (3.14%) | WUSTL | 1 (0.1%) |
| \$50,000-\$74,999 | 137 (13.0%) | | | YALE | 25 (2.4%) |
| \$75,000-\$99,999 | 115 (10.9%) | | | MSSM | 2 (0.2%) |
| \$100,000-\$199,999 | 116 (11.0%) | | | | |
| \$200,000 and greater | 21 (2.0%) | | | | |
| No answer | 135 (12.8%) |  |  |  |  |
| **Youth (N = 1,158)** |  |  |  |  |  |
| **Youth Gender** |  | ***n* (%)** |  |  |  |
| Male |  | 610 (52.7%) |  |  |  |
| Female |  | 547 (47.2%) |  |  |  |
| Refuse to answer |  | 1 (0.1%) |  |  |  |
| **Youth Nativity** |  |  |  |  |  |
| First-Generation |  | 96 (8.3%) |  |  |  |
| Second-Generation |  | 855 (73.8%) |  |  |  |
| Third-Generation |  | 207 (17.9%) |  |  |  |

*Note.* *n* = number of participants, % = percent of sample, *M* = mean, *SD* = standard deviation.

### Neuroimaging Data

Youth participants completed a baseline neuroimaging protocol that included structural MRI and rs-fMRI using high spatial and temporal resolution simultaneous multislice/multiband echo-planar (EPI) (46,47). For Siemens scanners, fMRI scan parameters were: 90×90 matrix, 60 slices, field of view=216×216, echo time / repetition time=30/ 800ms, flip angle=52°, 2.4mm isotropic resolution, and slice acceleration factor 6. The complete protocols for all vendors and sequences are provided by Casey et al. (46).

### Analyses

#### Latent Profile Analysis (LPA)

To empirically evaluate Berry’s model of acculturation by identifying distinct groups of caregivers based on their VIA scores (e.g., US and heritage subscales), we conducted an LPA in Mplus 8.7 with a Robust Maximum Likelihood (MLR) estimator and a sandwich covariance estimator to adjust the standard errors and account for the nesting of participants within site (48,49). A combination of fit statistics and substantive interpretability were used to decide on the number of profiles (50). First, a solution with *k* profiles was selected only if it provided a significantly better fit than a solution with *k*-1 profiles to balance parsimony and fit. This was determined using the Lo-Mendell-Rubin Adjusted Likelihood Ratio Test (LRT), which indicates the extent to which the −2 log likelihood value for a model with *k* profiles is significantly smaller than the corresponding value for a model with *k*-1 profiles. Second, entropy values and posterior probabilities of correct classification should be at least 0.70 (51). Third, to ensure stability of the profile solution, each profile had to represent more than 5% of the sample (52). Fourth, the profiles had to be conceptually and substantively different from one another; one profile could not simply be a variant of another profile.

#### Caregiver and Youth Mental Health

Next, we implemented the widely utilized classify-analyze approach (53) and saved profile membership and posterior probabilities for the championed profile model back into the dataset. This approach reduces uncertainty in profile classification and tends to not have the disadvantages associated with a one-step approach but can be biased when entropy is below 0.70 (54). Subsequently, we estimated a series of path models with acculturative orientation as a categorical predictor to determine if there were differences in terms of caregiver and youth mental health using the ASR and CBCL data, respectively. All subsequent path models were estimated in Mplus 8.7 (55) with an MRL estimator and a sandwich covariance estimator (48,49) to account for the nesting of participants within site. The Benjamini-Hochberg correction was applied to control for the false discovery rate (FDR) at 0.05 due to multiple comparisons (56). Covariates included caregiver education, identity, nativity, as well as youth gender and family income. Missing data were handled using full-information maximum likelihood estimation. Given the presence of siblings in the youth dataset, youth mental health models were estimated using multilevel modeling (MLM) to account for nesting of children (level 1) within family (level 2) and site (level 3).

#### Neuroimaging Preprocessing

MRI data were processed using fMRIPrep 21.0.0, a BIDS-App that automatically adapts a best-in-breed workflow, ensuring high-quality preprocessing with minimal manual intervention (57,58). Anatomical images were intensity-corrected, skull-stripped, segmented, and spatially normalized to a standard brain template in MNI space. Functional MRI preprocessing included motion correction, susceptibility distortion correction, and coregistration. Denoising was performed using AFNI’s (59) 3dTproject (60). Complete details of the MRI preprocessing and analyses are described in the SI.

#### Resting State fMRI Analyses

Resting state fMRI analyses were preregistered (https://osf.io/mkdw3/). We focused on five meta-analytically-defined regions of interest (ROIs) associated with self- and affiliation-related processing (61), including the (1) vmPFC, (2) left insula, (3) right insula, (4) left TPJ, and (5) right TPJ (**Fig. S1**). For each resting-state acquisition, voxelwise time series were extracted from each ROI using the unsmoothed, preprocessed, and denoised data via AFNI’s 3dmaskave. Averaged ROI time series were generated by calculating the mean voxel value for each time point across non-zero voxels respective to each region. For each ROI, we computed two measures of rs-fMRI activity: fractional amplitude of low-frequency fluctuations (fALFF) as a measure of local, spontaneous fluctuations during the resting state (62), and regional homogeneity (ReHo) as a measure of local BOLD signal coherence (63). In addition, adjacency matrices were constructed for each participant by computing the pairwise Pearson’s correlation coefficients between each pair of ROIs generating a 5×5 connectivity matrix with 10 unique regional pairs. Correlation values were Fisher *z*-transformed to provide a summary measure of pairwise functional connectivity.

Next, we estimated a series of path models to characterize potential differences across caregiver acculturation profiles in terms of rs-fMRI activity and connectivity. Three separate models were tested for: (1) spontaneous fluctuations using fALFF, (2) local signal coherence using ReHo, and (3) functional connectivity using z-transformed correlation coefficients. Path models were estimated in Mplus 8.7 controlling for caregiver education, identity, nativity, as well as youth gender and family income.

## RESULTS

### Latent Profile Analysis

As shown in **Table 2**, the 2-profile solution provided significantly better fit compared to the 1-profile solution [LRT=468.392, *p*<0.001]. Although entropy was higher, AIC/BIC were lower for the 3-profile solution, and the LRT test was trending towards significance [LRT=143.473, *p*=0.065], closer examination of the 3-profile solution indicated the additional profile was not conceptually and substantively different from one of the initial profiles and also only accounted for 4.16% of the sample (see SI). Thus, the 2-profile solution was advanced as the championed model.

**Table 2.** Latent Profile Analysis Model Comparisons.

| # of Profiles | AIC | BIC | Adj BIC | Entropy | Smallest | LRT (3) | p-Value |
| --- | --- | --- | --- | --- | --- | --- | --- |
| 2 | 5520.457 | 5555.199 | 5532.966 | 0.791 | 26.68% | 468.392 | <0.001 |
| 3 | 5376.117 | 5425.748 | 5393.987 | 0.833 | 4.16% | 143.472 | 0.065 |
| 4 | 5284.536 | 5349.057 | 5307.767 | 0.871 | 2.37% | 93.123 | 0.188 |

*Note.* AIC = Akaike Information Criteria, BIC = Bayesian Information Criteria, LRT = Lo-Mendell-Rubin Adjusted Likelihood Ratio Test.

**Table 3** presents mean *z*-scores for heritage and US cultural orientations across profiles, indicating how far each profile deviated from the total sample average. These *z*-scores can be interpreted as an effect size index. The first profile represented 73.3% of the sample (*n*=775) and was marked by high levels of both heritage and US cultural orientation. In line with Berry’s conceptualization, this model was labeled “Bicultural”. The second profile, labeled “Detached”, accounted for 26.7% of the sample (*n*=282) and was characterized by low levels of both heritage and US cultural orientation.

**Table 3.** Standardized Differences Across the 2-Profile Solution.

| Acculturation Dimension<br>% of Sample | Bicultural<br>73.3% |  | Detached<br>26.7% |  |
| --- | --- | --- | --- | --- |
|  | Mean | SD | Mean | SD |
| Heritage Orientation | 0.471 (7.790) | 0.557 (0.955) | -1.295 (4.760) | 0.781 (1.340) |
| US Orientation | 0.392 (7.606) | 0.681 (1.086) | -1.079 (5.260) | 0.941 (1.501) |

*Note.* Dimensions were standardized and should be interpreted as average z-scores indicating how far each profile deviates from the total sample average scores and from other profiles. Original unstandardized scores are reported in parentheses.

Next, given entropy was greater than 0.70, we utilized the classify-analyze approach (53) and saved profile membership and posterior probabilities back into the dataset. The average posterior probabilities were 0.96 and 0.89 for the bicultural and detached profiles, respectively. To ensure clearly defined class membership, we restricted profile assignment to those whose posterior probabilities were 0.70 or higher. Of the 1,057 total unique caregivers, 981 (92.81%) had posterior probabilities greater than 0.70, including 747 bicultural and 234 detached caregivers, and there were no significant differences between those with posterior probabilities below 0.70 (see SI). **Fig. 2** illustrates the *z*-scores for heritage and US cultural orientations across profiles. There were no significant differences between profiles in terms of caregiver identity, age, and generational status; family income; or youth gender and generational status. However, there was a significant difference in terms of education, such that caregivers with a bicultural orientation were overrepresented among higher educational attainment categories (**Table 4**).

**Figure 2.**
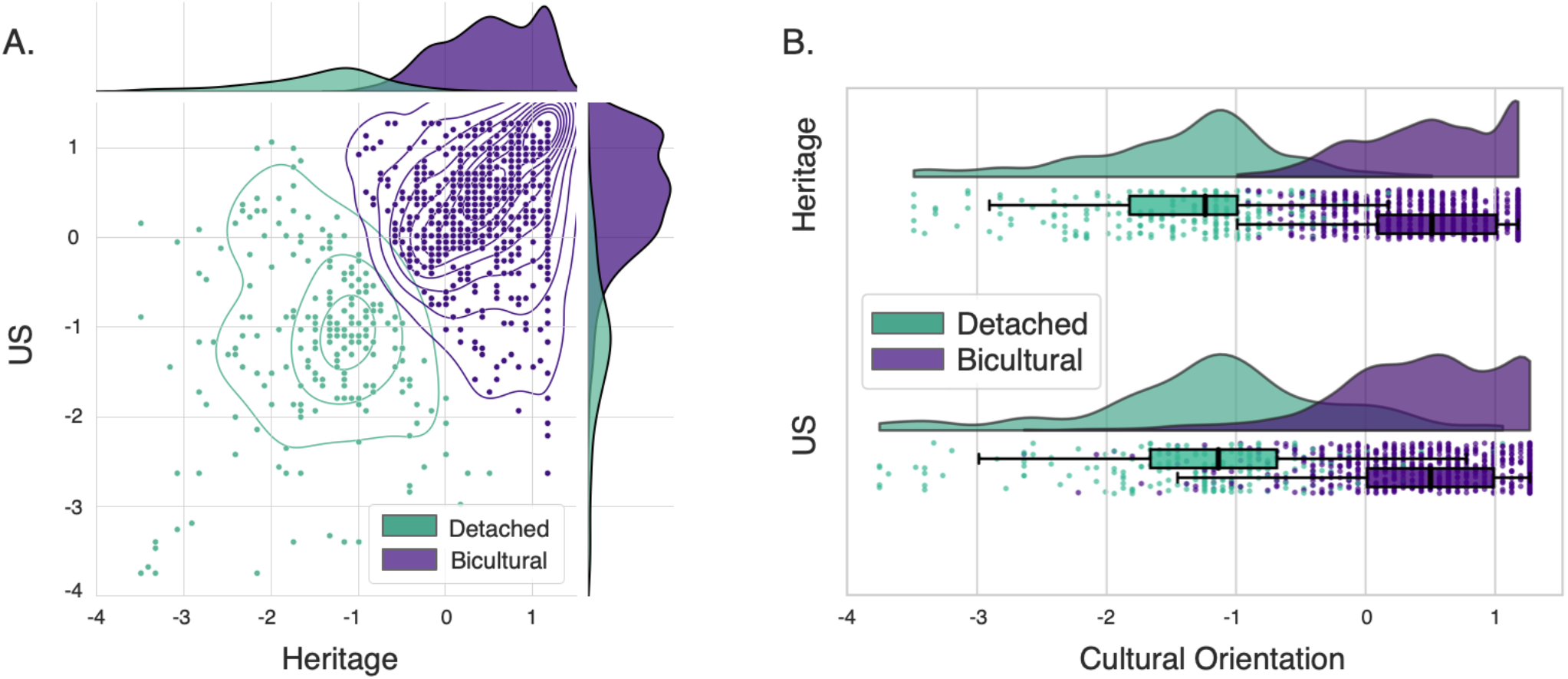
Heritage and US Cultural Orientations Across Profiles. LPA revealed two acculturative orientation profiles among Hispanic/Latinx caregivers in the ABCD Study. Cultural orientation subscale scores were normalized across participants and the resulting z-scored values are shown in: (A) a joint kernel density estimate plot (64) and (B) a raincloud plot (65). In both visualizations, the “Bicultural” profile (n=747) demonstrated high levels of both heritage and US cultural orientation (purple), while the “Detached” profile (n=234) exhibited very low levels of both heritage and US cultural orientation (green).

**Table 4.** Demographic Characteristics for Bicultural and Detached Groups.

|  | Total Caregiver Sample (N = 1,057) | Bicultural Caregivers (n = 747) | Detached Caregivers (n = 234) | Statistics |
| --- | --- | --- | --- | --- |
| **Caregiver Identity** | *n (%) or M (SD)* | *n (%) or M (SD)* | *n (%) or M (SD)* |  |
| Biological mother | 955 (90.4%) | 682 (91.3%) | 206 (88.0%) | $\chi^2(1) = 2.213$ |
| Biological father | 102 (9.6%) | 65 (8.7%) | 28 (12.0%) | $p = 0.137$ |
|  |  |  |  | *Cramér's V = 0.05* |
| **Caregiver Age** | 38.31 yrs (6.64 yrs) | 38.49 yrs (6.48 yrs) | 37.53 yrs (6.48 yrs) | $t(971) = -1.95$ |
| | | | | $p = 0.051$ |
|  |  |  |  | *Cohen's d = 0.15* |
| **Caregiver Nativity** |  |  |  |  |
| US born | 312 (29.5%) | 163 (21.8%) | 71 (30.3%) | $\chi^2(1) = 0.210$ |
| Foreign born | 745 (70.5%) | 532 (78.2%) | 163 (69.7%) | $p = 0.647$ |
|  |  |  |  | *Cramér's V = 0.02* |
| **Caregiver Education** |  |  |  |  |
| < High school diploma | 239 (22.6%) | 158 (21.2%) | 69 (29.5%) | $t(975) = -3.840$ |
| High school diploma/GED | 167 (15.8%) | 109 (14.6%) | 43 (18.4%) | $p < 0.001$ |
| Some college | 352 (33.3%) | 252 (33.7%) | 74 (31.6%) | *Cohen's d = 0.29* |
| Bachelor's degree | 172 (16.3%) | 127 (17.0%) | 31 (13.3%) |  |
| Postgraduate degree | 123 (11.6%) | 98 (13.1%) | 16 (6.8%) |  |
| No answer | 4 (0.4%) | 3 (0.4%) | 1 (0.4%) |  |
| **Household Income per Year (\$US)** | | | | |
| <\$5,000 | 49 (4.6%) | 37 (5.0%) | 7 (3.0%) | $t(853) = -0.436$ |
| \$5,000-\$11,999 | 65 (6.1%) | 48 (6.4%) | 14 (6.0%) | $p = 0.663$ |
| \$12,000-\$15,999 | 53 (5.0%) | 38 (5.1%) | 13 (5.6%) | *Cohen's d = 0.03* |
| \$16,000-\$24,999 | 103 (9.7%) | 66 (8.8%) | 34 (14.5%) | |
| \$25,000-\$34,999 | 126 (11.9%) | 89 (11.9%) | 23 (9.8%) | |
| \$35,000-\$49,999 | 137 (13.0%) | 103 (13.8%) | 26 (11.1%) | |
| \$50,000-\$74,999 | 137 (13.0%) | 96 (12.9%) | 30 (12.8%) | |
| \$75,000-\$99,999 | 115 (10.9%) | 82 (11.0%) | 26 (11.1%) | |
| \$100,000-\$199,999 | 116 (11.0%) | 84 (11.2%) | 21 (9.0%) | |
| \$200,000 and greater | 21 (2.0%) | 14 (1.9%) | 4 (1.7%) | |
| No answer | 135 (12.8%) | 90 (12.0%) | 36 (15.4%) |  |
|  | **Total Youth Sample (N = 1,158)** | **Youth w/ Bicultural Caregivers (n = 814)** | **Youth w/ Detached Caregivers (n = 261)** | **Statistics** |
| **Youth Gender** | *n (%)* | *n (%)* | *n (%)* |  |
| Male | 610 (52.7%) | 439 (53.9%) | 131 (50.2%) | $\chi^2(1) = 1.149$ |
| Female | 547 (47.2%) | 374 (46.0%) | 130 (49.8%) | $p = 0.284$ |
| No answer | 1 (0.1%) | 1 (0.1%) | 0 (0%) | *Cramér's V = 0.03* |
| **Youth Nativity** |  |  |  |  |
| First-Generation | 96 (8.3%) | 71 (8.7%) | 20 (7.7%) | $\chi^2(2) = 296$ |
| Second-Generation | 855 (73.8%) | 596 (73.2%) | 194 (74.3%) | $p = 0.863$ |
| Third-Generation | 207 (17.9%) | 147 (18.1%) | 47 (18.0%) | *Cramér's V = 0.02* |

*Note.* *n* = number of participants, % = percent of sample, *M* = mean, *SD* = standard deviation.

### Class Membership Effects: Caregiver and Youth Mental Health

Next, we estimated a path model with acculturative orientation included as a categorical predictor (0=detached, 1=bicultural) to determine whether there were significant differences across profiles in terms of caregiver mental health. Results indicated that being bicultural was associated with fewer symptoms of depression, avoidant, and inattentive behaviors; however, the association between caregivers’ acculturative orientation and inattentive behavior did not survive correction for multiple comparisons (**Table 5)**.

**Table 5.**
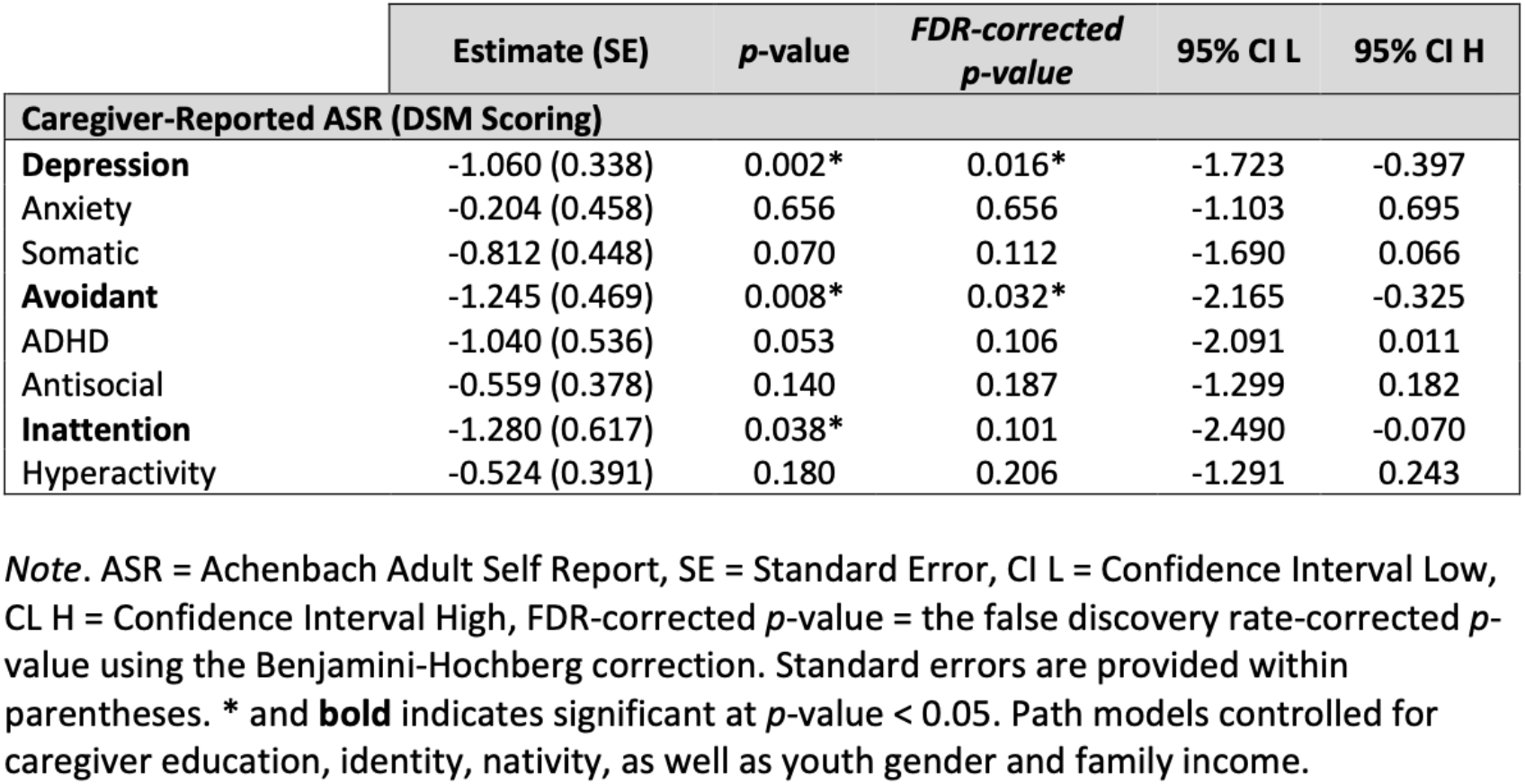
Class Membership Effects on Caregiver Mental Health.

|  | Estimate (SE) | *p*-value | *FDR-corrected*<br>*p-value* | 95% CI L | 95% CI H |
| --- | --- | --- | --- | --- | --- |
| **Caregiver-Reported ASR (DSM Scoring)** |  |  |  |  |  |
| **Depression** | -1.060 (0.338) | 0.002* | 0.016* | -1.723 | -0.397 |
| Anxiety | -0.204 (0.458) | 0.656 | 0.656 | -1.103 | 0.695 |
| Somatic | -0.812 (0.448) | 0.070 | 0.112 | -1.690 | 0.066 |
| **Avoidant** | -1.245 (0.469) | 0.008* | 0.032* | -2.165 | -0.325 |
| ADHD | -1.040 (0.536) | 0.053 | 0.106 | -2.091 | 0.011 |
| Antisocial | -0.559 (0.378) | 0.140 | 0.187 | -1.299 | 0.182 |
| **Inattention** | -1.280 (0.617) | 0.038* | 0.101 | -2.490 | -0.070 |
| Hyperactivity | -0.524 (0.391) | 0.180 | 0.206 | -1.291 | 0.243 |

*Note.* ASR = Achenbach Adult Self Report, SE = Standard Error, CI L = Confidence Interval Low, CI H = Confidence Interval High, FDR-corrected *p*-value = the false discovery rate-corrected *p*-value using the Benjamini-Hochberg correction. Standard errors are provided within parentheses. \* and **bold** indicates significant at *p*-value < 0.05. Path models controlled for caregiver education, identity, nativity, as well as youth gender and family income.

We examined whether there were significant differences between caregiver bicultural and detached profiles in terms of youth mental health. Children of caregivers with a bicultural orientation exhibited significantly fewer symptoms of depression and somatic complaints; however, neither of these findings survived the correction for multiple comparisons (**Table 6)**.

**Table 6.** Class Membership Effects on Youth Mental Health.

|  | Estimate (SE) | *p*-value | FDR-corrected<br>*p*-value | 95% CI L | 95% CI H |
| --- | --- | --- | --- | --- | --- |
| **Caregiver-Reported CBCL (DSM Scoring)** |  |  |  |  |  |
| **Depression** | -0.652 (0.318) | 0.040* | 0.120 | -1.276 | -0.029 |
| Anxiety | -0.386 (0.454) | 0.393 | 0.472 | -1.271 | 0.500 |
| **Somatic** | -0.850 (0.381) | 0.026* | 0.156 | -1.597 | -0.104 |
| ADHD | -0.371 (0.315) | 0.239 | 0.359 | -0.989 | 0.247 |
| Oppositional Defiant | -0.036 (0.248) | 0.886 | 0.886 | -0.522 | 0.451 |
| Conduct | -0.601 (0.410) | 0.142 | 0.284 | -1.404 | 0.202 |

*Note.* CBCL = Child Behavior Checklist .SE = Standard Error, CI L = Confidence Interval Low, CI H = Confidence Interval High, FDR-corrected *p*-value = the false discovery rate-corrected *p*-value using the Benjamini-Hochberg correction. Standard errors are provided within parentheses. \* and **bold** indicates significant at *p*-value < 0.05. Path models controlled for caregiver education, identity, nativity, as well as youth gender and family income.

### Class Membership Effects: Youth rs-fMRI Activity and Connectivity

Lastly, we examined whether there were significant differences between caregiver bicultural and detached profiles in terms of youth rs-fMRI activity and connectivity. With respect to spontaneous fluctuations during the resting state (**Table 7a)**, children of caregivers with a bicultural orientation exhibited greater fALFF values in the left insula; however, this finding did not survive correction for multiple comparisons. In terms of regional homogeneity (**Table 7b)**, children of caregivers with a bicultural orientation exhibited greater ReHo values (even after correction for multiple comparisons) also in the left insula. In terms of pairwise functional connectivity values (**Table 8)**, no significant differences in connectivity between the vmPFC, bilateral insula, and bilateral TPJ were observed among youth with bicultural vs. detached caregivers.

**Table 7.** Class Membership Effects on Youth rs-fMRI Activity.

|  | Estimate (SE) | *p*-value | FDR-corrected<br>*p*-value | 95% CI L | 95% CI H |
| --- | --- | --- | --- | --- | --- |
| **(a) fALFF** |  |  |  |  |  |
| vmPFC | 0.015 (0.017) | 0.400 | 0.800 | -0.020 | 0.049 |
| **Left Insula** | 0.021 (0.008) | 0.006* | 0.060 | 0.006 | 0.036 |
| Right Insula | 0.010 (0.013) | 0.418 | 0.597 | -0.014 | 0.035 |
| Left TPJ | -0.019 (0.022) | 0.394 | 0.985 | -0.063 | 0.025 |
| Right TPJ | -0.012 (0.015) | 0.417 | 0.695 | -0.041 | 0.017 |
| **(b) ReHo** |  |  |  |  |  |
| vmPFC | -0.004 (0.021) | 0.850 | 0.944 | -0.044 | 0.036 |
| **Left Insula** | 0.030 (0.011) | 0.007* | 0.035* | 0.008 | 0.051 |
| Right Insula | 0.025 (0.016) | 0.112 | 0.373 | -0.006 | 0.056 |
| Left TPJ | -0.015 (0.034) | 0.668 | 0.835 | -0.081 | 0.052 |
| Right TPJ | <.001 (0.030) | 0.995 | 0.995 | -0.059 | 0.059 |

*Note.* fALFF = fractional amplitude of low-frequency fluctuations, ReHo = regional homogeneity, vmPFC = ventromedial prefrontal cortex, TPJ = temporoparietal junction. SE = Standard Error, CI L = Confidence Interval Low, CI H = Confidence Interval High, FDR-corrected *p*-value = the false discovery rate-corrected *p*-value using the Benjamini-Hochberg correction. Standard errors are provided within parentheses. \* and **bold** indicates significant at *p*-value < 0.05. Path models controlled for caregiver education, identity, nativity, as well as youth gender and family income.

**Table 8.**
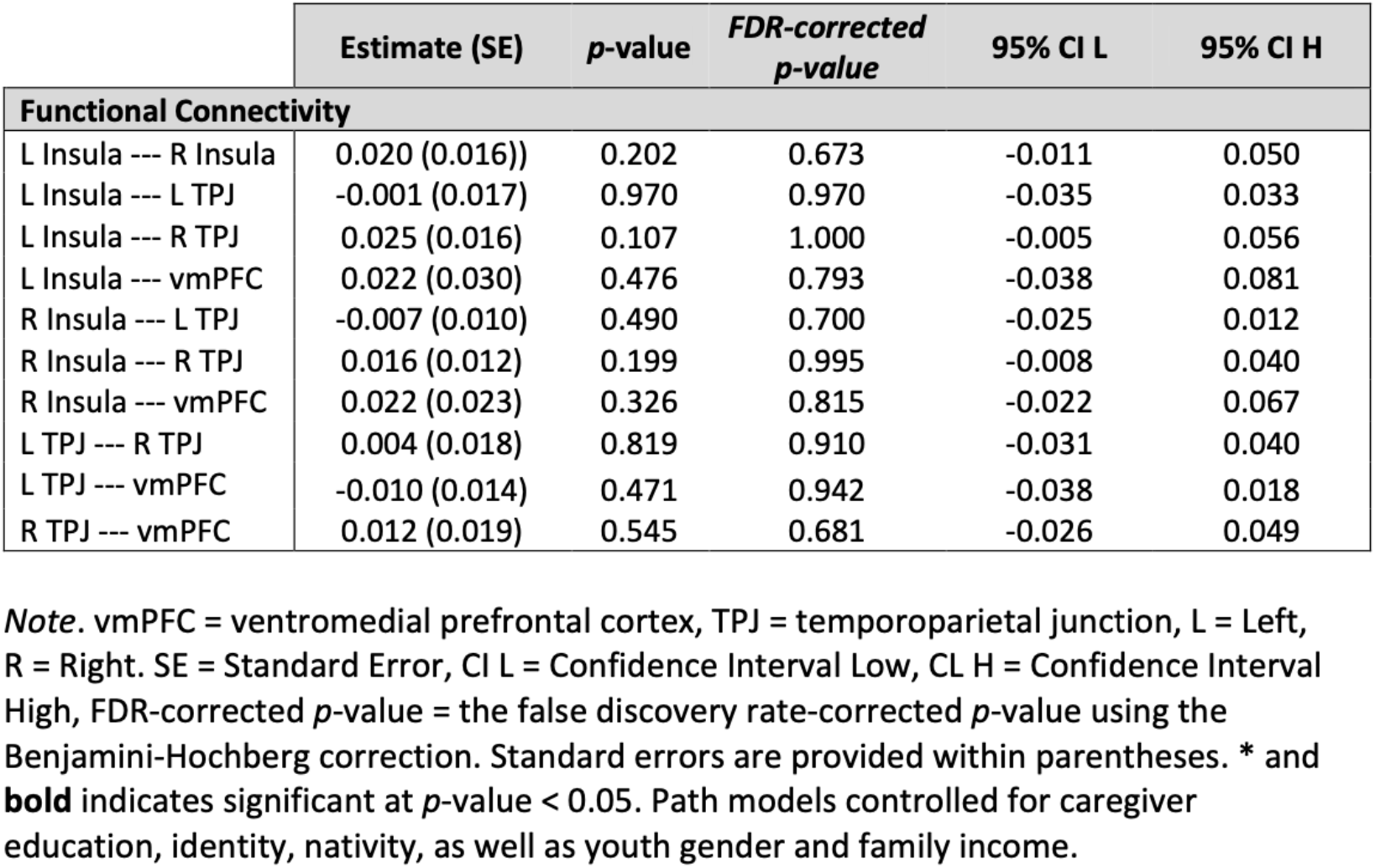
Class Membership Effects on Youth rs-fMRI Connectivity.

|  | Estimate (SE) | *p*-value | *FDR-corrected*<br>*p-value* | 95% CI L | 95% CI H |
| --- | --- | --- | --- | --- | --- |
| **Functional Connectivity** |  |  |  |  |  |
| L Insula --- R Insula | 0.020 (0.016) | 0.202 | 0.673 | -0.011 | 0.050 |
| L Insula --- L TPJ | -0.001 (0.017) | 0.970 | 0.970 | -0.035 | 0.033 |
| L Insula --- R TPJ | 0.025 (0.016) | 0.107 | 1.000 | -0.005 | 0.056 |
| L Insula --- vmPFC | 0.022 (0.030) | 0.476 | 0.793 | -0.038 | 0.081 |
| R Insula --- L TPJ | -0.007 (0.010) | 0.490 | 0.700 | -0.025 | 0.012 |
| R Insula --- R TPJ | 0.016 (0.012) | 0.199 | 0.995 | -0.008 | 0.040 |
| R Insula --- vmPFC | 0.022 (0.023) | 0.326 | 0.815 | -0.022 | 0.067 |
| L TPJ --- R TPJ | 0.004 (0.018) | 0.819 | 0.910 | -0.031 | 0.040 |
| L TPJ --- vmPFC | -0.010 (0.014) | 0.471 | 0.942 | -0.038 | 0.018 |
| R TPJ --- vmPFC | 0.012 (0.019) | 0.545 | 0.681 | -0.026 | 0.049 |

*Note.* vmPFC = ventromedial prefrontal cortex, TPJ = temporoparietal junction, L = Left, R = Right. SE = Standard Error, CI L = Confidence Interval Low, CI H = Confidence Interval High, FDR-corrected *p*-value = the false discovery rate-corrected *p*-value using the Benjamini-Hochberg correction. Standard errors are provided within parentheses. \* and **bold** indicates significant at *p*-value < 0.05. Path models controlled for caregiver education, identity, nativity, as well as youth gender and family income.

## DISCUSSION

Acculturation data from Hispanic/Latinx caregivers in the ABCD Study were analyzed and yielded two caregiver profiles. “Bicultural” caregivers (n=747) endorsed high levels of both heritage and US cultural orientation, while “detached” (n=234) caregivers endorsed very low levels of heritage and US cultural orientation. Theory (8–10) and prior experimental work (11,13,14) have identified two additional profiles, “assimilated” (high US and low heritage) and “separated” (low US and high heritage), which were surprisingly not observed. It is possible that separated caregivers were less likely to enroll their children in the ABCD Study due to mistrust, documentation concerns, and/or reluctance to participate in a high-profile, national, longitudinal study (66). Lack of the assimilated profile may have stemmed from using the VIA, which captures bidimensional acculturation specifically in terms of cultural practices. Given that acculturation extends across multiple domains (1), future research should utilize measures of heritage and US identification and values to better capture nuances in cultural orientations. That said, it is worth noting past research has not always consistently identified assimilated (21,67–69) or separated profiles (67,70) among adult populations. As a whole, in contrast to results among adolescents and young adults (37), these findings may indicate decreased variability in cultural orientation among adults. Overall, utilization of a data-driven approach allowed for the testing of competing models underlying the data, yielding more ecologically valid findings that reflect the lived experiences of ABCD Hispanic/Latinx families.

Comparison of ABCD bicultural and detached profiles indicated that biculturalism was associated with more positive mental health outcomes in agreement with a wealth of previous acculturation research (10,18,19,71,72). However, these prior studies are characterized by the use of individual rating inventories that separately measure symptoms of a single disorder (e.g., depression). The current work represents the first time the ASR has been utilized to study acculturative orientations, allowing for a more comprehensive evaluation across multiple symptoms and disorders. We observed an interesting pattern of behavioral problems among detached caregivers, including increased avoidant behaviors and symptoms of depression and inattention. When placed in context of need and motivation for relationships with others, this pattern suggests a diminished approach for and enjoyment of social interactions. It is possible that individuals with a detached orientation experience overall low social affiliation that influences their connectedness to other people and broader social structures. Future work will need to address the extent to which detached orientations may reflect low behavioral activation system (BAS) (73) sensitivity or anhedonia and low social approach (74–76), resulting in reduced effort to obtain rewards and/or low sensitivity to rewards. Such links would represent significant progress in establishing clinical phenotypes of acculturation that have been previously related to outcomes such as depression (17,20), substance use (72,77), and suicide risk (71).

From a family systems perspective, current findings within the ABCD sample indicate potential downstream effects among youth as a result of caregiver acculturation. The results suggest significant, and likely complex, differences in caregiving environments as a result of caregiver health, parenting, family functioning, that collectively and dynamically influences fetal through adolescent development. Emerging patterns of increased symptoms of depression and somatic complaints among 9-10-year-old youth with detached caregivers were observed, thus identifying a subset of Hispanic/Latinx youth that are at higher risk for adverse outcomes at the beginning of adolescence, a critical socioemotional neurodevelopmental period wherein psychopathology often emerges (78,79). Importantly, these results are derived from caregiver-reported measures, which can be influenced by the depression-distortion bias (80), and perhaps provide somewhat limited insight into youth perspectives on acculturation and mental health. Future analyses of subsequent ABCD time points should incorporate youth-reported measures. Furthermore, the longitudinal design of the ABCD Study provides a unique opportunity to continue following these at-risk Hispanic/Latinx youth, with the aim of more fully understanding divergent trajectories sensitive to stressful family dynamics that arise during adolescence when Hispanic/Latinx youth are developing their own cultural values, beliefs, and practices (30,81). Such analyses would be enhanced by the inclusion of mediating variables related to family conflict, prosocial behaviors, and parenting. Given the comprehensive youth assessments included in the ABCD Study, future research also offers the opportunity to directly assess links between adolescent acculturation and low BAS sensitivity or anhedonic phenotypes.

In terms of neural differences, youth of bicultural caregivers exhibited greater rs-fMRI activity (both fALFF and ReHo) in the left insula, potentially indicating increased metabolic rate of glucose and oxygen (82); these results are aligned with studies showing the insula is associated with somatic and depressive symptoms (83,84) and psychopathology broadly (85–87). The insula is a complex, multifaceted structure (88). If indeed detached caregivers transmit their acculturative orientation directly to their children, and if detached orientations are linked to anhedonia and/or altered reward behaviors, then it is possible that dysregulated insula function is a central neurobiological mechanism of interest, given its prominent role in reward processing (89,90). Finally, we note that while differences in rs-fMRI activity (i.e., fALFF and ReHo) were observed, significant effects were not found for long-range connectivity differences. Future work should examine neurodevelopmental changes across adolescence to determine if localized, corticolimbic brain effects among Hispanic/Latinx youth at ABCD baseline are ultimately translated into long-range connectivity differences.

In conclusion, these findings indicate that acculturation is an important factor relevant to ABCD Hispanic/Latinx caregivers’ mental health, as well as the mental health and resting-state insula activity of their children. This work demonstrates that the ABCD multisite design and demographically diverse sample offer an opportunity to study participant groups that have been historically underrepresented in neuroimaging research. Further, the current work intentionally does not compare Hispanic vs. non-Hispanic participants, which can implicitly provide support for a deficits-based framework. Instead, we emphasize the diversity of Hispanic/Latinx families in the US who have diverse family dynamics, life experiences, and health-related outcomes. Moreover, analysis of a subset of the ABCD sample creates a space to focus on Hispanic/Latinx culture in a way that is not centered around non-Hispanic, majority experiences. ABCD’s robust culture and environment protocol was thoughtfully developed and provided the measures that made this work feasible. Additional population-based, neuroimaging studies incorporating other culturally-relevant measures of the social and structural determinants of health are urgently needed. Such work will allow for a more complete understanding of neurobiological processes of risk and resilience among individuals from underrepresented, minoritized groups that experience health-related disparities as a consequence of their racial, ethnic, sexual, or gender identity.

## Data Availability

Data used in the preparation of this article were obtained from the Adolescent Brain Cognitive Development^SM^ (ABCD) Study (https://abcdstudy.org), held in the NIMH Data Archive (NDA). This is a multisite, longitudinal study designed to recruit more than 10,000 children age 9-10 and follow them over 10 years into early adulthood. The ABCD Study® is supported by the National Institutes of Health and additional federal partners under award numbers U01DA041048, U01DA050989, U01DA051016, U01DA041022, U01DA051018, U01DA051037, U01DA050987, U01DA041174, U01DA041106, U01DA041117, U01DA041028, U01DA041134, U01DA050988, U01DA051039, U01DA041156, U01DA041025, U01DA041120, U01DA051038, U01DA041148, U01DA041093, U01DA041089, U24DA041123, U24DA041147. A full list of supporters is available at https://abcdstudy.org/federal-partners.html. A listing of participating sites and a complete listing of the study investigators can be found at https://abcdstudy.org/consortium_members/. ABCD consortium investigators designed and implemented the study and/or provided data but did not necessarily participate in the analysis or writing of this report. This manuscript reflects the views of the authors and may not reflect the opinions or views of the NIH or ABCD consortium investigators.

The ABCD data repository grows and changes over time. The ABCD data used in this report came from NDA ABCD Release 3.0 (DOI: 10.15154/1520591). An NDA Study was created to associate the analyses reported in this study with the underlying ABCD Study data (DOI: 10.15154/1527751).

## Code and Materials Availability

Additional information and resources are available on a project page for this study at the Open Science Framework (OSF) (https://osf.io/mkdw3/). The code is available in two GitHub repositories, including one for analysis of the VIA data (https://github.com/NBCLab/abcd-hispanic-via) and one for the fMRI preprocessing and analyses (https://github.com/NBCLab/abcd_fmriprep-analysis). ROI and connectivity maps are available in NeuroVault (https://identifiers.org/neurovault.collection:1276). High-resolution figures are available via FigShare (https://figshare.com/account/home#/projects/143997).

## Author Contributions

AM and ARL conceived and designed the project. AM, JAP, MCR, WH, and ARL analyzed data. JAP, MCR, TS, JSF, and KLB contributed scripts and pipelines. WH, ASD, and MTS provided statistical support and JWP, RPL, LMU, CAG, SWH, MS, MRG, RG provided support for analysis of ABCD measures and results interpretation. AM, JAP, MCR, and ARL wrote the paper and all authors contributed to the revisions and approved the final version.

## Footnotes

1 Although Berry (1997) originally used the term “Integration”, the dual endorsement of heritage and US cultures has been increasingly referred to as “Biculturalism” (1). Moreover, although Berry used the term “Marginalized” to represent detachment from both heritage and US cultures, we use the term “Detached” as a better representation of individuals’ lack of connection to either cultural stream.

