## Supplemental Information for "Acculturative orientations among Hispanic/Latinx caregivers in the ABCD Study: Associations with caregiver and youth mental health and youth brain function"

### SUPPLEMENTAL INTRODUCTION

Large, population-based neuroscience studies offer opportunities to investigate research questions among more diverse and representative samples (Falk et al., 2013). However, this diversity may often implicitly serve as an encouragement for neuroimaging researchers to stratify samples and examine brain-based group differences on the basis of ethnicity, gender, race, sex, or other relevant demographic information. Such analyses often unintentionally provide support for a biological deficits-based framework, which argues that lower rates of achievement and/or poorer life outcomes among minoritized groups are the result of "biological determinism." Responsible and ethical data analyses should strive to prevent stigmatizing and marginalizing research through consideration of potentially harmful impacts research could have (Laird, 2021; Nketia et al., 2021; Saragosa-Harris et al., 2021; Simmons et al., 2021). Importantly, it is critical that researchers consider the impact of factors related to social determinants of health (WHO, 2022). The goal of this study was to demonstrate the importance of considering cultural variables, with a focus on the exemplar domain of acculturation, which has been linked to a variety of outcomes, including alcohol use (Lui and Zamboanga, 2018), cigarette smoking (Meca et al., 2017), and suicide risk (Meca et al., 2022).

### SUPPLEMENTAL METHODS

#### Participants

Participants were selected from the ABCD Study, the largest long-term study of brain development and child health in the United States (US) (Volkow et al., 2018). Approximately 11,800 youth aged 9.00 to 10.99 years old were enrolled in the ABCD Study across 21 sites in the US (Garavan et al., 2018). Participants within the ABCD Study were recruited through geographically, demographically, and socioeconomically diverse school systems using epidemiologically informed methods to enroll a population-based, demographically diverse sample (Compton et al., 2019). The ABCD Study was approved by the Institutional Review Board (IRB) at each study site and centralized IRB approval was provided by the University of California San Diego. All child participants provided informed assent to participate while caregivers (i.e., parent or legal guardian) gave informed consent. Additional information regarding recruitment and assessment procedures have been extensively reported elsewhere (Dick et al., 2021; Volkow et al., 2018). Data from the ABCD Study are made available by the NIMH Data Archive (NDA; https://nda.nih.gov) and the current study utilized data from the ABCD Curated Annual Release 3.0.

#### Vancouver Index of Acculturation

Caregivers completed the Vancouver Index of Acculturation (VIA) in English or Spanish. The VIA is a 16-item bidimensional measure with subscales that separately measure heritage and US acculturation (Ryder et al., 2000). Items addressed a range of topics, including traditions, social activities, media, cultural values, and behavioral preferences (**Table S1**). Initial assessment of VIA data at ABCD baseline indicated high internal consistency across the heritage ($\alpha$=0.92) and US ($\alpha$=0.90) subscales, with early data suggesting higher VIA subscale scores for both heritage and US cultures among families at lower risk compared to those at higher risk for adolescent substance use (Zucker et al., 2018).

**Table S1. Caregiver-Reported Vancouver Index of Acculturation (VIA) Items for the ABCD Study.**

| Heritage Items | Mainstream Items |
|---|---|
| **H1**: I often participate in my heritage cultural traditions. | **US1**: I often participate in mainstream American cultural traditions. |
| **H2**: I enjoy social activities with people from the same heritage culture as myself. | **US2**: I enjoy social activities with typical American people. |
| **H3**: I am comfortable interacting with people of the same heritage culture as myself. | **US3**: I am comfortable interacting with typical American people. |
| **H4**: I enjoy entertainment (e.g., movies, music) from my heritage culture. | **US4**: I enjoy typical American entertainment (e.g., movies, music). |
| **H5**: I often behave in ways that are typical of my heritage culture. | **US5**: I often behave in ways that are typically American. |
| **H6**: It is important for me to maintain or develop the practices of my heritage culture. | **US6**: It is important for me to maintain or develop American mainstream cultural practices. |
| **H7**: I believe in the values of my heritage culture. | **US7**: I believe in mainstream American values. |
| **H8**: I am interested in having friends from my heritage culture. | **US8**: I am interested in having typical American friends. |

*Note*. The VIA is a 16-item bidimensional measure with subscales that separately measure heritage and US acculturation. Caregivers were asked to provide their heritage culture as an open-ended item with specific examples provided as prompts (e.g., "*Asian*", "*Black/African American*", "*Hispanic*", "*Native American*", "Jewish"); those who did not identify a heritage culture were told not to complete the VIA.

#### Hispanic/Latinx Sample Selection

A total of 11,878 ABCD participants were recruited at baseline. Data for the present analyses were downloaded from NDA for 2,411 participants who completed their baseline assessment for the ABCD Study and responded "*Yes*" to "*Do you consider the child Hispanic/Latino/Latina?*" (`demo_ethn_v2`). We filtered participant datasets to include only caregivers who: (1) completed the VIA, (2) referenced Hispanic/Latinx culture when completing the VIA, (3) and were either first- or second-generation immigrants from Latin America. Generational status was determined based on data regarding caregivers' and maternal and paternal grandparents' nativity. Further details on family immigration history are not available in the ABCD dataset, preventing differentiation of later-generation immigrants. That is, it is not possible to ascertain if a caregiver is a third-, fourth-, or ninth-generation immigrant. Prior studies have documented a general increase in US cultural orientation and decrease in heritage cultural orientation with each successive generation (Yoon et al., 2020). Inclusion of later-generation Hispanic/Latinx caregivers in the present study would introduce substantial variability. Thus, to reduce heterogeneity, we restricted the sample to include only caregivers who were either first- or second-generation immigrants, and subsequently, youth who were either first-, second-, or third-generation immigrants.

Given uncertainty with assessing generational status among non-biological caregivers, 87 non-biological caregivers were removed from the dataset. Subsequently, 8 families were removed due to lack of information on reporting child and/or caregiver nativity. An additional 944 caregivers were removed after indicating that neither their children, themselves, and their own parents were born in a Latin

American country. Next, because the VIA represents the primary variable of interest, of the remaining 1,372 participants, 37 were removed after not responding to a single item of the VIA. Participants with missing data did not differ from those with complete data with regards to caregiver identity [$\chi 2$ (1) = 0.175, $p$ = 0.676], caregiver generational status [$\chi 2$ (1) = 0.801, $p$ = 0.371], education [$\chi 2$ (4) = 6.241, $p$ = 0.182], family income [$t$(1178) = -0.307, $p$ = 0.759], youth gender [$\chi 2$ (1) = 0.060, $p$ = 0.807], and youth generational status [$\chi 2$ (2) = 0.304, $p$ = 0.859]. Additionally, because the VIA focuses on culture broadly, it was important to make sure participants were responding to questions based on their ethnic culture. As such, 111 were removed after not selecting a particular cultural group and an additional 66 were removed after referring to a non-Hispanic/Latinx cultural group (e.g., religious affiliation), respectively.

The subsequent sample thus consisted of 1,057 caregivers (mean age = 38.31 years, *SD* = 6.64 years; 90.4% mothers and 9.6% fathers, 70.5% foreign born) and 1,158 children (52.7% male, 91.7% US born). In nearly all cases (*n* = 203 siblings), the same caregiver reported either at the same time point or at a subsequent time point for their multiple children participating in the study. To avoid nesting within-time among caregivers, we selected the earliest data point from the caregiver for a given family to ensure all children in that same family unit solely had the same caregiver point of data. For the remaining 8 participants, we randomly selected one of the two caregivers to represent the family and paired data with all siblings within that family. **Fig. S1** provides details on the analytic sample.

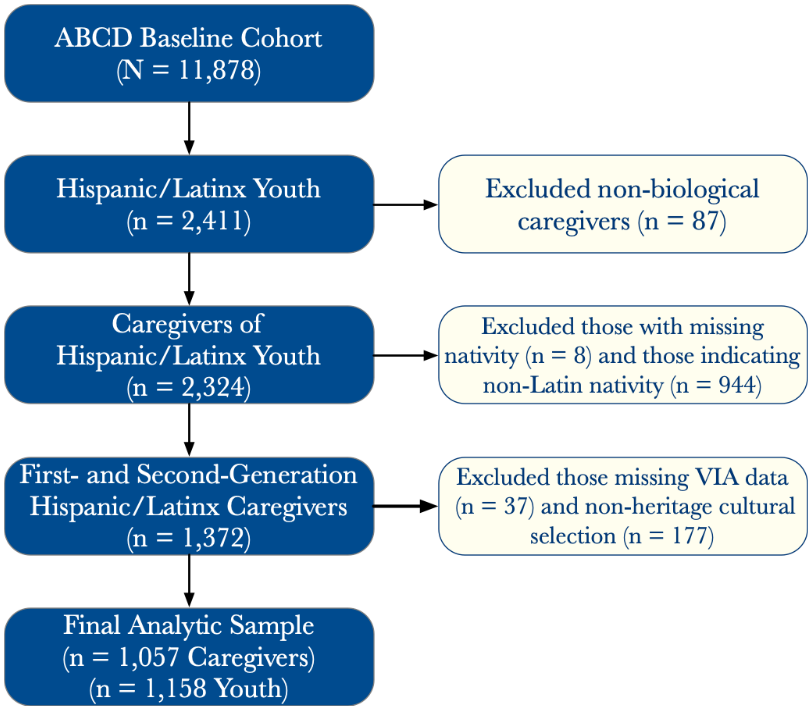


**Figure S1. Inclusion and exclusion criteria for the analytic sample selection.**

#### Neuroimaging Data

Youth participants completed a baseline neuroimaging protocol that included structural magnetic resonance imaging (MRI), as well as resting-state functional MRI using high spatial and temporal resolution simultaneous multislice/multiband echo-planar (EPI) (Casey et al., 2018; Hagler et al., 2019). For Siemens scanners, scan parameters were: 90 × 90 matrix, 60 slices, field of view (FOV) = 216 × 216, echo time (TE)/repetition time (TR) (ms) = 30/800, flip angle = 52°, 2.4 mm isotropic resolution, and slice acceleration factor 6. The complete protocols for all vendors and sequences are provided by Casey et al. (Casey et al., 2018).

##### Download and Conversion to BIDS

ABCD data are housed on Amazon Web Service's (AWS) Simple Storage Solution (S3). S3 links can be downloaded using AWS's command line tools (`aws-cli`). The resulting file associated with each link corresponds to a compressed folder containing the DICOM images for a specific participant, session, and acquisition (i.e., T1-weighted, T2-weighted, BOLD, etc.).

The *abcd-dicom2bids* repository (https://github.com/DCAN-Labs/abcd-dicom2bids) is a robust package that downloads ABCD magnetic resonance imaging (MRI) data and converts the DICOM data organization to fit the Brain Imaging Data Structure (BIDS) specification (Gorgolewski et al., 2016). Aside from software dependencies, an additional requirement is that one must download the ABCD Quality Control Spreadsheet (`abcd_fastqc01.csv`), which can be downloaded from the NIMH Data Archive with an approved Data Use Certification (DUC) that provides access to ABCD Study data. This quality control spreadsheet contains the AWS S3 links to each participant's modality-specific acquisition and is annotated with a 1 or a 0 depending on whether DAIC quality control annotators found the acquisition to be free of artifacts and suitable for analysis. The ABCD Release 3.0 Quality Control spreadsheet was downloaded on May 03, 2021.

While the *abcd-dicom2bids* repository is an immensely useful and valuable resource, we noted some drawbacks to using it in our lab's ABCD workflows. Thus, we chose to generate a modified version of the repository for the current study. First, the current release of *abcd-dicom2bids* only supports downloading event-related files in text format. The modified version allows for downloading task-related event information files stored in either text (`.txt`) or comma-separated value (`.csv`) format. Second, the modified version permits download of specific functional tasks, rather than only having the option to download all functional tasks, which is more of an enhancement than a drawback. Third, in some instances, the *abcd-dicom2bids* repository deleted erroneously identified DICOMs for each TR acquired by Philips scanners. This ultimately resulted in the failure to convert the DICOMs to BIDS format. Fourth, and most relevant to the current study, the wide range of dependencies needed for operation is a limiting factor when attempting to use this package on a High Performance Cluster (HPC) where administrative privileges may not be available. In this instance, provided the HPC has Singularity or Docker installed, using a container or image is advantageous because no administrative privileges are needed for software installation.

Thus, we generated a Docker image https://hub.docker.com/repository/docker/julioaperaza/abcddicom2bids of our modified version of the *abcd-dicom2bids* repository https://github.com/JulioAPeraza/abcd-dicom2bids, which was then converted to a Singularity image on FIU's High Performance Cluster (HPC).

#### Neuroimaging Preprocessing

MRI data were processed using `fMRIPrep` 21.0.0, a BIDS-App that automatically adapts a best-in-breed workflow, ensuring high-quality preprocessing with minimal manual intervention (Esteban et al., 2020, 2019).

#### Anatomical Data Preprocessing

T1-weighted (T1w) images were corrected for intensity non-uniformity (INU) with `N4BiasFieldCorrection` (Tustison et al., 2010), distributed with `ANTs` 2.3.3 (Avants et al., 2008) (RRID:SCR_004757) and used as a T1w-reference throughout the workflow. The T1w-reference was then skull-stripped with a Nipype implementation of the `antsBrainExtraction.sh` workflow (from `ANTs`), using `OASIS30ANTs` as the target template. Brain tissue segmentation of cerebrospinal fluid (CSF), white-matter (WM), and gray-matter (GM) was performed on the brain-extracted T1w using `fast` (`FSL` 6.0.5.1:57b01774, RRID:SCR_002823) (Zhang et al., 2001). Brain surfaces were reconstructed using `recon-all` (`FreeSurfer` 6.0.1, RRID:SCR_001847) (Dale et al., 1999) and the brain mask estimated previously was refined with a custom variation of the method to reconcile ANTs-derived and FreeSurfer-derived segmentations of the cortical gray-matter of `Mindboggle` (RRID:SCR_002438) (Klein et al., 2017). Volume-based spatial normalization to one standard space (*MNI152NLin2009cAsym*) was performed through nonlinear registration with `antsRegistration` (`ANTs` 2.3.3), using brain-extracted versions of both the T1w reference and T1w template. The following template was selected for spatial normalization: ICBM 152 Nonlinear Asymmetrical template version 2009c [(Fonov et al., 2009), RRID:SCR_008796; `TemplateFlow` ID: *MNI152NLin2009cAsym*].

#### Functional Data Preprocessing

Each participant's dataset contained 1-4 runs (i.e., acquisitions) of resting state functional magnetic resonance imaging (rs-fMRI) data. For each of the BOLD runs found per subject, the following preprocessing was performed. First, a reference volume and its skull-stripped version were generated using a custom methodology of `fMRIPrep`.

Head motion parameters with respect to the BOLD reference (transformation matrices, and six corresponding rotation and translation parameters) were estimated before any spatiotemporal filtering using `mcflirt` (`FSL` 6.0.5.1:57b01774) (Jenkinson et al., 2002). The estimated fieldmap was then aligned with rigid-registration to the target EPI (echo-planar imaging) reference run. The field coefficients were mapped on to the reference EPI using the transform. The BOLD reference was then co-registered to the T1w reference using `bbregister` (`FreeSurfer`), which implements boundary-based registration (Greve and Fischl, 2009). Co-registration was configured with six degrees of freedom. Several confounding time series were calculated based on the preprocessed BOLD: framewise displacement (FD), DVARS, and three region-wise global signals. FD was computed using two formulations following Power et al. (i.e., absolute sum of relative motions (Power et al., 2014)) and Jenkinson et al. (i.e., relative root mean square displacement between affines (Jenkinson et al., 2002)). FD and DVARS were calculated for each functional run, both using their implementations in `Nipype` (following the definitions by Power et al. (Power et al., 2014)). The three global signals were extracted within the CSF, the WM, and the whole-brain masks. Additionally, a set of physiological regressors were extracted to allow for component-based noise correction (`CompCor` (Behzadi et al., 2007)). Principal components were estimated after high-pass filtering the preprocessed BOLD time series (using a discrete cosine filter with 128s cut-off) for the two `CompCor` variants: temporal (*tCompCor*) and anatomical (*aCompCor*). *tCompCor* components were then calculated from the top 2% variable voxels within the brain mask. For *aCompCor*, three probabilistic masks (CSF, WM, and combined CSF+WM) were generated in anatomical space. The implementation differs from that of (Behzadi et al., 2007) in that instead of eroding the masks by 2 pixels on BOLD space, the *aCompCor* masks subtracted a mask of pixels that likely contained a volume fraction of GM. This mask was obtained by dilating a GM mask extracted from the `FreeSurfer`'s *aseg* segmentation, and it ensured components were not extracted from voxels containing a minimal fraction of GM. Finally, these masks were resampled into BOLD space and binarized by thresholding at 0.99 (as in the original implementation). Components were also calculated separately within the WM and CSF masks. For each `CompCor` decomposition, the *k* components with the largest singular values were retained, such that the retained components' time series were sufficient to explain 50 percent of variance across the nuisance mask (CSF, WM, combined, or temporal). The remaining components were dropped from consideration. The head motion estimates calculated in the correction step were also placed within the corresponding confounds file. The confound time series derived from head motion estimates and global signals were expanded with the inclusion of temporal derivatives and quadratic terms for each (Satterthwaite et al., 2013). All resamplings can be performed with a single interpolation step by composing all the pertinent transformations (i.e., head-motion transform matrices, susceptibility distortion correction when available, and co-registrations to anatomical and output spaces). Gridded (volumetric) resamplings were performed using `antsApplyTransforms` (`ANTs`), configured with Lanczos interpolation to minimize the smoothing effects of other kernels (Lanczos, 1964). Non-gridded (surface) resamplings were performed using `mri_vol2surf` (`FreeSurfer`).

Many internal operations of `fMRIPrep` use `Nilearn` 0.6.2 (Abraham et al., 2014) (RRID:SCR_001362), mostly within the functional processing workflow. For more details of the pipeline, see the section corresponding to workflows in fMRIPrep's documentation: https://fmriprep.org/en/latest/workflows.html.

AFNI's `3dTproject` was used to perform simultaneous denoising of nuisance regressors and bandpass filtering. Nuisance regressors included global signal regression (GSR), ten *aCompCor* components (i.e., 5 CSF, 5 WM) (Muschelli et al., 2014), the six motion parameters and their derivatives, and TRs acquired during MRI stabilization (i.e., non-steady state), as determined by the ABCD Consortium in accordance with scanner manufacturer recommendations (Siemens/Philips: 8 TRs; GE: 5 TRs). A 0.01 to 0.1 Hz bandpass filter was also applied. Functional volumes with framewise displacement (FD) greater than 0.35mm, and the time points immediately preceding and following, were removed after nuisance signal regression. As recommended by Hagler et al. (Hagler et al., 2019), BOLD runs with fewer than 100 usable time points (out of a total of 375 acquired) were excluded from further analysis.

`3dTproject` was performed three times, separately, on `fMRIPrep` preprocessed data: (1) without bandpass filtering and without smoothing, (2) with bandpass filtering and without smoothing, and (3) with bandpass filtering and with 6-mm FWHM blur applied after regression. This step yielded three datasets. The first dataset was used to estimate the fALFF (described below), and the other two datasets, which differed by the application of the 6-mm blur, were used for extracting an ROI time series from the unsmoothed dataset that was used in the deconvolution of the smoothed dataset.

`MRIQC` 0.16.1 (Esteban et al., 2017), a BIDS-App that computes quality control (QC) measures, was used to compute resting state QC measures of signal-to-noise ratio (SNR), temporal SNR, mean framewise displacement (FD), and ghost-to-signal ratio in the x- and y-directions, and entropy focus criterion.

#### Neuroimaging Analyses

##### Meta-Analytic Regions of Interest (ROIs)

Regions of interest were defined using a consensus-based approach that integrated findings across multiple prior neuroimaging meta-analyses. First, identified neural regions that would be sensitive to processes associated with identity development; thus, we targeted regions previously associated with self-related processing and social affiliations. To this end, we drew from a recent meta-analysis of social processing (Pintos Lobo et al., 2022) that was structured according to the NIMH's RDoC framework (Insel, 2014). Comparisons across neuroimaging studies revealed that the RDoC Social Processes domain construct of *Perception and Understanding of Self* was selectively associated with the ventromedial prefrontal cortex (vmPFC) (cluster centroid of [*-6, 48, -8*]) and left temporoparietal junction (TPJ) (cluster centroid of [*-44, -56, 24*]), while the *Affiliation and Attachment* construct was selectively associated with the left insula (cluster centroid of [*-40, 18, 0*]) and left TPJ (cluster centroid of [*-44, -56, 28*]).

Second, we aimed to incorporate the delineation of ROI boundaries by drawing from robust, multimodal meta-analytic results that optimally represent functionally dissociable regions. We assessed spatial overlap between self- and affiliation-related clusters (Pintos Lobo et al., 2022) and previous large-scale, connectivity-based, and meta-analytic parcellations of these regions. Using this approach, we selected the dorsoanterior cluster of the central ventromedial frontal lobe identified by Chase et al. (i.e., Cluster 3) (Chase et al., 2020) (**Fig. S2A**; purple) and the bilateral dorsoanterior insula cluster identified by Chang et al. (Chang et al., 2013) (**Fig. S2A**; blue and green). For the TPJ ROI, we note that the *left* TPJ was observed to be relevant for both self- and affiliation-related processing (Pintos Lobo et al., 2022); however, meta-analytic assessment of the TPJ has focused on the *right* TPJ (Bzdok et al., 2013), given a lack of consensus regarding the left TPJ's role in social cognition. Thus, we selected the larger left TPJ cluster associated with self processing from the Pintos Lobo et al. meta-analysis (**Fig. S2B**; pink) and the right posterior TPJ region from the Bzdok meta-analysis (**Fig. S2B**; cyan). This ROI selection process allowed us to delineate five consensus-based, meta-analytically derived regions of interest likely to be associated with identity development for the (1) vmPFC, (2) left insula, (3) right insula, (4) left TPJ, and (5) right TPJ. Bilateral regions (e.g., left and right TPJ) were treated as separate ROIs. Altogether, this was a principled approach for selecting well-defined seeds representing multiple complex, heterogeneous brain regions.

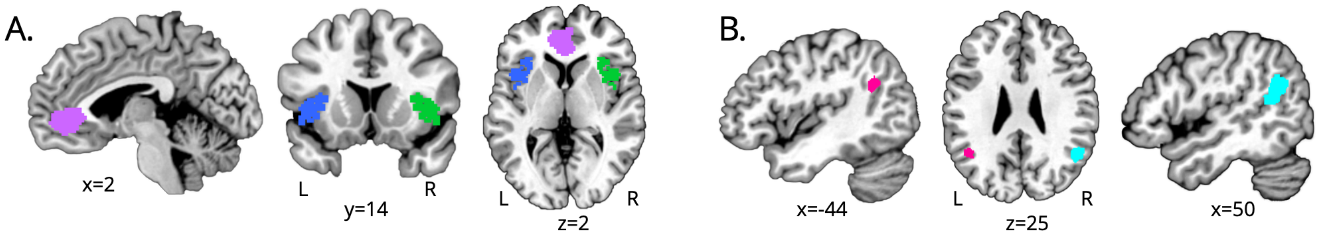


**Figure S2. Meta-Analytic Seeds for vmPFC, Insula, and TJP.** (A) The dorsoanterior cluster of the central ventromedial frontal lobe (purple) (Chase et al., 2020) was selected as the vmPFC ROI, while the left (blue) and right (green) dorsoanterior insula clusters (Chang et al., 2013) were selected as the left and right insula ROIs. (B) The left TPJ ROI (pink) was selected from a meta-analysis of self-related processing (Pintos Lobo et al., 2022), while the right posterior TPJ region (cyan) was selected from a meta-analysis of the right TPJ (Bzdok et al., 2013).

#### Resting State fMRI Analyses

Resting state fMRI analyses were preregistered (https://osf.io/mkdw3/). For each resting-state acquisition, voxelwise time series were extracted for each of the five ROIs from the unsmoothed, preprocessed, and denoised data using AFNI's `3dmaskave`. Averaged ROI time series were generated by calculating the mean voxel value for each time point across non-zero voxels respective to each region. For each ROI, we computed two different rs-fMRI measures of local activity and one measure of long-range functional connectivity.

##### rs-fMRI Local Spontaneous Activity

First, we used the fractional amplitude of low-frequency fluctuations (fALFF) as a measure of local, spontaneous fluctuations of the brain during the resting state. fALFF values were calculated at every voxel in each ROI by computing the ratio of the sum of amplitudes across the total frequency range (0.01-0.08 Hz) against the available frequency band (Zou et al., 2008) and averaging across voxels in each ROI. fALFF was estimated on the time series without bandpass filtering and without smoothing using AFNI's `3dLombScargle` and `3dAmpToRSFC` programs (Press and Rybicki, 1989). The combination of these two programs allow the estimation of rsFC parameters from time series with nonconstant sampling (e.g., censored data). Second, we used regional homogeneity (ReHo) as a measure of local BOLD signal coherence (Zang et al., 2004). ReHo was calculated on the unsmoothed dataset using AFNI's `ReHo` by computing the Kendall's coefficient of concordance of a voxel and its nearest neighbors (i.e., 27 voxels) and averaging across voxels in each ROI (Taylor and Saad, 2013). fALFF and ReHo values were computed for each of the five regions.

##### rs-fMRI Functional Connectivity

Adjacency matrices were constructed for each participant by computing the pairwise Pearson's correlation coefficient between each pair of ROIs, yielding a 5x5 connectivity matrix. Correlation values were Fisher *z*-transformed to provide a summary measure of pairwise functional connectivity.

##### Class Membership Effects: Youth rs-fMRI Activity and Connectivity

Next, we estimated a series of path models to determine if there were differences across extracted caregiver acculturation profiles in terms of rs-fMRI activity and connectivity. Three separate models were tested for (1) spontaneous fluctuations (i.e., fALFF), (2) local signal coherence (i.e., ReHo), and (iii) functional connectivity (i.e., *z*-transformed correlation coefficients). Path models were estimated in `Mplus` 8.7 (Muthén and Muthén, 1997) with a Robust Maximum Likelihood (MLR) estimator and a sandwich covariance estimator (Freedman, 2006; Kauermann and Carroll, 2001) to adjust the standard errors and account for the nesting of participants within site. Caregiver identity, nativity, and education, as well as youth gender and family income were included as covariates. Missing data were handled using full-information maximum likelihood estimation. Given the presence of siblings in the youth dataset, youth models were estimated using multilevel modeling (MLM) to account for nesting of children (level 1) within family (level 2) and site (level 3).

### SUPPLEMENTAL RESULTS

#### Latent Profile Analysis

The 2-profile solution was advanced as the championed model and the results of this model are reported in the manuscript. However, the 3-profile solution had higher levels of entropy and lower levels of AIC and BIC than the 2-profile solution and the LRT was trending towards significance. Our decision to advance the 2-profile solution over the 3-profile solution was guided by a number of reasons. First, although trending, the LRT was non-significant. It is worth noting when the sample is very large, traditional class enumeration procedures will reward models that continuously recover a finer structure with more classes, even if not always be of substantive interest (Curran and Bauer, 2021). As a result, it may be prudent in these situations to de-emphasizing information criteria based on concerns associated with overfitting a model. Second, the third profile identified only accounted for 4.16% of the sample.

Lastly, and most critical, the third extracted profile was not conceptually and substantively different from either of the initial profiles in terms of mean z-scores for heritage and US cultural orientations (**Table S2**). Taken together, the 2-profile solution was advanced as the championed model.

**Table S2. Standardized Differences Across the 3-Profile Solution.**

| Acculturation Dimension<br>% of Sample | Profile 1<br>66.3% | Profile 2<br>29.5% | Profile 3<br>4.2% |
|---|---|---|---|
| Heritage Orientation | 0.566 | -0.868 | -2.532 |
| US Orientation | 0.447 | -0.693 | -1.941 |

*Note*. Dimensions were standardized and should be interpreted as average z-scores indicating how far each profile deviates from the total sample average scores and from other profiles.

#### LPA Class Assignment

Following the LPA, profile membership and posterior probabilities were saved back into the dataset. To ensure clearly defined class membership, we restricted assignment to profiles to those whose posterior probabilities were 0.70 or higher. Of the 1,057 total unique caregiver cases, 981 of participants (92.81%) had posterior probabilities greater than 0.70. This included 747 bicultural and 234 detached caregivers. The remaining caregivers were excluded from all subsequent analyses. It is worth noting, those with posterior probabilities under 0.70 did not significantly differ from those with posterior probabilities above 0.70 in the context of caregiver identity [$\chi 2$ (1) = 0.945, $p$ = 0.331], youth gender [$\chi 2$ (1) = 1.349, $p$ = 0.245], caregiver generation [$\chi 2$ (1) = 0.867, $p$ = 0.352], youth generation [$\chi 2$ (3) = 2.201, $p$ = 0.532], caregiver education [$t$(920) = -1.137, $p$ = 0.256], and family income [$t$(1051) = -0.869, $p$ = 0.385].

### SUPPLEMENTAL ANALYSES

#### LPA Using Constituent VIA Items

A strength of LPA is its capacity to handle multivariate data and classify individuals across a number of indicators. That said, although the strength of LPA is in working with multivariate data, the goal of LPA, similar to other person-centered approaches, is to classify individuals regardless of the number of indicators that are utilized for this purpose. In the context of our specific study, our approach is paralleling the broader acculturation literature that has often utilized higher-ordered factors capturing heritage and US cultural orientation. Not only is this approach consistent with Berry's (1980) conceptualization (Berry, 1980), but it is aligned with how the Vancouver Index of Acculturation is typically utilized (Ryder et al., 2000; Testa et al., 2019) and consistent with a plethora of studies that have sought to experimentally validate Berry's bidimensional acculturation mode (Des Rosiers et al., 2013; Fox et al., 2013; Meca et al., 2018; Ren et al., 2021; Schwartz and Zamboanga, 2008; Yan et al., 2021).

To determine the consistency of our results when compared to alternative analytic approaches, we conducted an LPA using all of the VIA items. With this approach, the 2-profile solution once again emerged as the championed model over the 3-profile solution (**Table S3**). Moreover, the 2-profile solution largely was equivalent to a solution characterized by bicultural (68.59%) and detached (31.41%) profiles across heritage and US orientation items (**Table S4**).

**Table S3. Multivariate Latent Profile Analysis Model Comparisons.**

| # of Profiles | AIC | BIC | Adj BIC | Entropy | Smallest | LRT (17) | p-Value |
|---|---|---|---|---|---|---|---|
| 2 | 42589.53 | 42832.73 | 42677.10 | 0.943 | 31.41% | 0.0804 | 0.0804 |
| 3 | 41178.98 | 41506.55 | 41296.92 | 0.910 | 14.19% | 0.5062 | 0.5062 |
| 4 | 39991.47 | 40403.41 | 40139.79 | 0.933 | 8.89% | 0.5755 | 0.5755 |

*Note*. AIC = Akaike Information Criteria, BIC = Bayesian Information Criteria, LRT = Lo-Mendell-Rubin Adjusted Likelihood Ratio Test.

**Table S4. Standardized Differences Across the 2-Profile Solution.**

| Acculturation Dimension | Bicultural | Detached |
|---|---|---|
| % of Sample | 68.6% | 31.4% |
| **Heritage Orientation** | | |
| VIA_H1 | -0.980 | 0.452 |
| VIA_H2 | -1.209 | 0.558 |
| VIA_H3 | -1.180 | 0.544 |
| VIA_H4 | -0.865 | 0.399 |
| VIA_H5 | -0.933 | 0.431 |
| VIA_H6 | -1.184 | 0.546 |
| VIA_H7 | -1.295 | 0.598 |
| VIA_H8 | -1.121 | 0.517 |
| **US Orientation** | | |
| VIA_US1 | -0.797 | 0.368 |
| VIA_US2 | -1.059 | 0.489 |
| VIA_US3 | -0.951 | 0.439 |
| VIA_US4 | -0.893 | 0.412 |
| VIA_US5 | -0.615 | 0.284 |
| VIA_US6 | -1.076 | 0.497 |
| VIA_US7 | -1.072 | 0.495 |
| VIA_US8 | -1.157 | 0.534 |

*Note*. Dimensions were standardized and should be interpreted as average z-scores indicating how far each profile deviates from the total sample average scores and from other profiles.

Hagler, D.J., Hatton, S., Cornejo, M.D., Makowski, C., Fair, D.A., Dick, A.S., Sutherland, M.T., Casey, B.J., Barch, D.M., Harms, M.P., Watts, R., Bjork, J.M., Garavan, H.P., Hilmer, L., Pung, C.J., Sicat, C.S., Kuperman, J., Bartsch, H., Xue, F., Heitzeg, M.M., Laird, A.R., Trinh, T.T., Gonzalez, R., Tapert, S.F., Riedel, M.C., Squeglia, L.M., Hyde, L.W., Rosenberg, M.D., Earl, E.A., Howlett, K.D., Baker, F.C., Soules, M., Diaz, J., de Leon, O.R., Thompson, W.K., Neale, M.C., Herting, M., Sowell, E.R., Alvarez, R.P., Hawes, S.W., Sanchez, M., Bodurka, J., Breslin, F.J., Morris, A.S., Paulus, M.P., Simmons, W.K., Polimeni, J.R., van der Kouwe, A., Nencka, A.S., Gray, K.M., Pierpaoli, C., Matochik, J.A., Noronha, A., Aklin, W.M., Conway, K., Glantz, M., Hoffman, E., Little, R., Lopez, M., Pariyadath, V., Weiss, S.R., Wolff-Hughes, D.L., DelCarmen-Wiggins, R.,

Feldstein Ewing, S.W., Miranda-Dominguez, O., Nagel, B.J., Perrone, A.J., Sturgeon, D.T., Goldstone, A., Pfefferbaum, A., Pohl, K.M., Prouty, D., Uban, K., Bookheimer, S.Y., Dapretto, M., Galvan, A., Bagot, K., Giedd, J., Infante, M.A., Jacobus, J., Patrick, K., Shilling, P.D., Desikan, R., Li, Y., Sugrue, L., Banich, M.T., Friedman, N., Hewitt, J.K., Hopfer, C., Sakai, J., Tanabe, J., Cottler, L.B., Nixon, S.J., Chang, L., Cloak, C., Ernst, T., Reeves, G., Kennedy, D.N., Heeringa, S., Peltier, S., Schulenberg, J., Sripada, C., Zucker, R.A., Iacono, W.G., Luciana, M., Calabro, F.J., Clark, D.B., Lewis, D.A., Luna, B., Schirda, C., Brima, T., Foxe, J.J., Freedman, E.G., Mruzek, D.W., Mason, M.J., Huber, R., McGlade, E., Prescot, A., Renshaw, P.F., Yurgelun-Todd, D.A., Allgaier, N.A., Dumas, J.A., Ivanova, M., Potter, A., Florsheim, P., Larson, C., Lisdahl, K., Charness, M.E., Fuemmeler, B., Hettema, J.M., Maes, H.H., Steinberg, J., Anokhin, A.P., Glaser, P., Heath, A.C., Madden, P.A., Baskin-Sommers, A., Constable, R.T., Grant, S.J., Dowling, G.J., Brown, S.A., Jernigan, T.L., Dale, A.M., 2019. Image processing and analysis methods for the Adolescent Brain Cognitive Development Study. Neuroimage 202, 116091. https://doi.org/10.1016/j.neuroimage.2019.116091

Insel, T.R., 2014. The NIMH Research Domain Criteria (RDoC) Project: precision medicine for psychiatry. Am J Psychiatry 171, 395–397. https://doi.org/10.1176/appi.ajp.2014.14020138

Jenkinson, M., Bannister, P., Brady, M., Smith, S., 2002. Improved optimization for the robust and accurate linear registration and motion correction of brain images. Neuroimage 17, 825–841. https://doi.org/10.1016/s1053-8119(02)91132-8

Kauermann, G., Carroll, R.J., 2001. A Note on the Efficiency of Sandwich Covariance Matrix Estimation. Journal of the American Statistical Association 96, 1387–1396. https://doi.org/10.1198/016214501753382309

Klein, A., Ghosh, S.S., Bao, F.S., Giard, J., Häme, Y., Stavsky, E., Lee, N., Rossa, B., Reuter, M., Neto, E.C., Keshavan, A., 2017. Mindboggling morphometry of human brains. PLOS Computational Biology 13, e1005350. https://doi.org/10.1371/journal.pcbi.1005350

Laird, A.R., 2021. Large, open datasets for human connectomics research: Considerations for reproducible and responsible data use. Neuroimage 244, 118579. https://doi.org/10.1016/j.neuroimage.2021.118579

Lanczos, C., 1964. Evaluation of Noisy Data. Journal of the Society for Industrial and Applied Mathematics Series B Numerical Analysis 1, 76–85. https://doi.org/10.1137/0701007

Lui, P.P., Zamboanga, B.L., 2018. Acculturation and alcohol use among Asian Americans: A meta-analytic review. Psychol Addict Behav 32, 173–186. https://doi.org/10.1037/adb0000340

Meca, A., Cobb, C.L., Schwartz, S.J., Szabó, Á., Moise, R., Zamboanga, B.L., Lee, T.K., Klimstra, T.A., Soares, M.H., Ritchie, R., Stephens, D.P., 2018. Exploring Individual Differences in the Relationship Between Cultural Identity Processes and Well-Being. Emerging Adulthood 2167696818817168. https://doi.org/10.1177/2167696818817168

Meca, A., Polanco-Roman, L., Cowan, I., Jimenez, A., 2022. Understanding Suicide-Related Risk in Immigrant and Ethnic and Racial Minority Youth Through an Ecological and Developmental Context, in: Miranda, R., Jeglic, E.L. (Eds.), Handbook of Youth Suicide Prevention: Integrating Research into Practice. Springer International Publishing, Cham, pp. 115–135. https://doi.org/10.1007/978-3-030-82465-5_8

Meca, A., Reinke, L.G., Scheier, L.M., 2017. Acculturation and Tobacco/Illicit Drug Use in Hispanic Youth [WWW Document]. The Oxford Handbook of Acculturation and Health. https://doi.org/10.1093/oxfordhb/9780190215217.013.20

Muschelli, J., Nebel, M.B., Caffo, B.S., Barber, A.D., Pekar, J.J., Mostofsky, S.H., 2014. Reduction of motion-related artifacts in resting state fMRI using aCompCor. Neuroimage 96, 22–35. https://doi.org/10.1016/j.neuroimage.2014.03.028

Muthén, L.K., Muthén, B., 1997. Mplus User's Guide. Muthén & Muthén, Los Angeles, CA.

Nketia, J., Amso, D., Brito, N.H., 2021. Towards a more inclusive and equitable developmental cognitive neuroscience. Dev Cogn Neurosci 52, 101014. https://doi.org/10.1016/j.dcn.2021.101014

Pintos Lobo, R., Bottenhorn, K.L., Riedel, M.C., Toma, A.I., Hare, M.M., Smith, D.D., Moor, A.C., Cowan, I.K., Valdes, J.A., Bartley, J.E., Salo, T., Boeving, E.R., Pankey, B., Sutherland, M.T., Musser, E.D., Laird, A.R., 2022. Neural systems underlying RDoC social constructs: An activation likelihood estimation meta-analysis. Neurosci Biobehav Rev 144, 104971. https://doi.org/10.1016/j.neubiorev.2022.104971

Power, J.D., Mitra, A., Laumann, T.O., Snyder, A.Z., Schlaggar, B.L., Petersen, S.E., 2014. Methods to detect, characterize, and remove motion artifact in resting state fMRI. Neuroimage 84. https://doi.org/10.1016/j.neuroimage.2013.08.048

Press, W.H., Rybicki, G.B., 1989. Fast Algorithm for Spectral Analysis of Unevenly Sampled Data. The Astrophysical Journal 338, 277. https://doi.org/10.1086/167197

Ren, H., Cheah, C.S.L., Tahseen, M., Zhou, N., 2021. A person-centered examination of acculturation and psychological functioning among Chinese and Korean immigrant mothers in the United States. Cultural Diversity and Ethnic Minority Psychology 27, 95–106. https://doi.org/10.1037/cdp0000430

Ryder, A.G., Alden, L.E., Paulhus, D.L., 2000. Is acculturation unidimensional or bidimensional? A head-to-head comparison in the prediction of personality, self-identity, and adjustment. J Pers Soc Psychol 79, 49–65.

Saragosa-Harris, N., Chaku, N., MacSweeney, N., Williamson, V.G., Scheuplein, M., Feola, B., Cardenas-Iniguez, C., Demir-Lira, E., McNeilly, E.A., Huffman, L.G., Whitmore, L., Michalska, K.J., Damme, K.S., Rakesh, D., Mills, K., 2021. A practical guide for researchers and reviewers using the ABCD Study and other large longitudinal datasets. https://doi.org/10.31234/osf.io/aqckx

Satterthwaite, T.D., Elliott, M.A., Gerraty, R.T., Ruparel, K., Loughead, J., Calkins, M.E., Eickhoff, S.B., Hakonarson, H., Gur, R.C., Gur, R.E., Wolf, D.H., 2013. An improved framework for confound regression and filtering for control of motion artifact in the preprocessing of resting-state functional connectivity data. NeuroImage 64, 240–256. https://doi.org/10.1016/j.neuroimage.2012.08.052

Schwartz, S.J., Zamboanga, B.L., 2008. Testing Berry's model of acculturation: a confirmatory latent class approach. Cultur Divers Ethnic Minor Psychol 14, 275–285. https://doi.org/10.1037/a0012818

Simmons, C., Conley, M.I., Gee, D.G., Baskin-Sommers, A., Barch, D.M., Hoffman, E.A., Huber, R.S., Iacono, W.G., Nagel, B.J., Palmer, C.E., Sheth, C.S., Sowell, E.R., Thompson, W.K., Casey, B.J., 2021. Responsible Use of Open-Access Developmental Data: The Adolescent Brain Cognitive Development (ABCD) Study. Psychol Sci 32, 866–870. https://doi.org/10.1177/09567976211003564

Taylor, P.A., Saad, Z.S., 2013. FATCAT: (An Efficient) Functional And Tractographic Connectivity Analysis Toolbox. Brain Connectivity 3, 523–535. https://doi.org/10.1089/brain.2013.0154

Testa, S., Doucerain, M.M., Miglietta, A., Jurcik, T., Ryder, A.G., Gattino, S., 2019. The Vancouver Index of Acculturation (VIA): New evidence on dimensionality and measurement invariance across two cultural settings. International Journal of Intercultural Relations 71, 60–71. https://doi.org/10.1016/j.ijintrel.2019.04.001

Tustison, N.J., Avants, B.B., Cook, P.A., Zheng, Y., Egan, A., Yushkevich, P.A., Gee, J.C., 2010. N4ITK: Improved N3 Bias Correction. IEEE Transactions on Medical Imaging 29, 1310–1320. https://doi.org/10.1109/TMI.2010.2046908

Volkow, N.D., Koob, G.F., Croyle, R.T., Bianchi, D.W., Gordon, J.A., Koroshetz, W.J., Pérez-Stable, E.J.,

Riley, W.T., Bloch, M.H., Conway, K., Deeds, B.G., Dowling, G.J., Grant, S., Howlett, K.D., Matochik, J.A., Morgan, G.D., Murray, M.M., Noronha, A., Spong, C.Y., Wargo, E.M., Warren, K.R., Weiss, S.R.B., 2018. The conception of the ABCD study: From substance use to a broad NIH collaboration. Dev Cogn Neurosci 32, 4–7. https://doi.org/10.1016/j.dcn.2017.10.002

WHO, W.H.O., 2022. Social Determinants of Health [WWW Document]. URL https://www.who.int/westernpacific/health-topics/social-determinants-of-health (accessed 3.11.22).

Yan, J., Sim, L., Schwartz, S.J., Shen, Y., Parra-Medina, D., Kim, S.Y., 2021. Longitudinal profiles of acculturation and developmental outcomes among Mexican-origin adolescents from immigrant families. New Dir Child Adolesc Dev 2021, 205–225. https://doi.org/10.1002/cad.20396

Yoon, E., Cabirou, L., Galvin, S., Hill, L., Daskalova, P., Bhang, C., Ahmad Mustaffa, E., Dao, A., Thomas, K., Baltazar, B., 2020. A Meta-Analysis of Acculturation and Enculturation: Bilinear, Multidimensional, and Context-Dependent Processes. The Counseling Psychologist 48, 342–376. https://doi.org/10.1177/0011000019898583

Zang, Y., Jiang, T., Lu, Y., He, Y., Tian, L., 2004. Regional homogeneity approach to fMRI data analysis. Neuroimage 22, 394–400. https://doi.org/10.1016/j.neuroimage.2003.12.030

Zhang, Y., Brady, M., Smith, S., 2001. Segmentation of brain MR images through a hidden Markov random field model and the expectation-maximization algorithm. IEEE Transactions on Medical Imaging 20, 45–57. https://doi.org/10.1109/42.906424

Zou, Q.-H., Zhu, C.-Z., Yang, Y., Zuo, X.-N., Long, X.-Y., Cao, Q.-J., Wang, Y.-F., Zang, Y.-F., 2008. An improved approach to detection of amplitude of low-frequency fluctuation (ALFF) for resting-state fMRI: fractional ALFF. J. Neurosci. Methods 172, 137–141. https://doi.org/10.1016/j.jneumeth.2008.04.012

Zucker, R.A., Gonzalez, R., Feldstein Ewing, S.W., Paulus, M.P., Arroyo, J., Fuligni, A., Morris, A.S., Sanchez, M., Wills, T., 2018. Assessment of culture and environment in the Adolescent Brain and Cognitive Development Study: Rationale, description of measures, and early data. Dev Cogn Neurosci 32, 107–120. https://doi.org/10.1016/j.dcn.2018.03.004
